# Cell cycle dynamics regulate H3K27 and H3K9 histone modifications

**DOI:** 10.1101/2025.08.18.670808

**Authors:** Liyne Nogay, Ananthakrishnan Vijayakumar Maya, Lara Heckmann, Francesco Cardamone, Isabelle Grass, Aakriti Singh, Anna Frey, Laurin Ernst, Nicola Iovino, Anne-Kathrin Classen

**Affiliations:** Faculty of Biology, University of Freiburg, Freiburg, Germany; International Max Planck Research School for Epigenetics, Biophysics and Metabolism, Freiburg, Germany; Max Planck Institute of Immunobiology and Epigenetics, Freiburg, Germany; CIBSS Centre for Integrative Biological Signalling Studies, University of Freiburg, Freiburg, Germany; Spemann Graduate School of Biology and Medicine (SGBM), University of Freiburg Germany

## Abstract

Cell cycle progression presents a fundamental challenge to genome integrity, particularly due to the need to reestablish post-translational histone modifications (PTMs) following DNA replication. Although proliferative and differentiating tissues exhibit markedly different cell cycle dynamics, how these differences shape the histone modification landscape in vivo remains largely unexplored. Here, we show that levels of H3K27ac, H3K27me3, and H3K9me3 are tightly linked to cell cycle dynamics in the *Drosophila* wing imaginal disc. We demonstrate that both physiological and pathological elongation of the cell cycle led to an accumulation of H3K9me3 and H3K27me3, whereas cell cycle acceleration reduces their levels. In contrast, H3K27ac exhibits the opposite pattern: levels decrease in arrested cells and increase with faster cycling. Genome-wide CUT&Tag analysis reveals that these changes predominantly affect genomic loci already modified in normally proliferating tissue. Importantly, the regulation of methylation levels at H3K9 and H3K27 is not solely mediated by the cell cycle machinery but reflects a metabolically guided process in which the rate of methylation is coupled to the rate of cell proliferation through metabolic activity, including signaling via the Insulin/PI3K/Akt pathway. Our study thus reveals key principles for understanding histone methylation in proliferating, senescent, and differentiating cells. In contrast, H3K27 acetylation is regulated through a distinct, cell cycle-coupled mechanism. We find that CBP/Nejire-mediated acetylation of H3K27 peaks during S-phase and is reversed by HDAC1, as cells exit replication. Disruption of this acetylation cycle leads to replication stress and a G2 cell cycle arrest via a DNA damage checkpoint. Notably, this genome-protective function of CBP/Nejire depends specifically on acetylation of the H3K27 residue itself, revealing a novel role for H3K27ac beyond its well-established function in transcriptional activation. Together, our findings establish a robust link between cell cycle progression and histone modification dynamics, highlighting the necessity of maintaining balanced PTM levels under varying proliferative states. These insights have broad implications for our understanding of development, aging, and tumor growth.

## INTRODUCTION

The cell cycle and the regulation of chromatin need to be tightly coordinated preserve epigenetic integrity. Specifically, progression through S-phase poses several challenges: chromatin must become transiently accessible for DNA replication, pre-existing post-translationally modified histones must be evenly distributed between daughter genomes and newly incorporated histones must acquire appropriate modifications to compensate for the semi-conservative dilution of modifications. These demands are complicated by the fact that cell cycle progression and dynamics differ drastically across biological contexts, ranging from rapid divisions in embryonic and regenerating tissues to complete arrest in terminally differentiated cells. How chromatin states are maintained across diverse proliferative tissue environments *in vivo* is still insufficiently understood.

In actively proliferating tissues, entry into the cell cycle at the G1/S transition, and progression through subsequent cycle phases, are tightly regulated. This regulation is mediated by Cyclins and Cyclin-dependent kinases (CDKs), as well as retinoblastoma (Rb) proteins and E2F transcription factors. Throughout the cell cycle, specific checkpoints respond to growth factors or cell size to modulate the rate of proliferation, as well as to DNA damage or chromosome misalignment to maintain the integrity of the genome (Matthews et al., 2022, Tapon et al., 2001, Lee and Orr-Weaver, 2003). As cells enter stages of differentiation, quiescence or senescence, they typically withdraw from active cycling by either prolonging their G1 phase or entering a stable cell cycle arrest known as G0. This transition away from proliferation often involves cyclin-dependent kinase (CDK) inhibitors, such as p21 and p27, which are intricately linked to gene expression programs that guide cell fate determination and differentiation (Buttitta and Edgar, 2007, Huang et al., 2022, de Morree and Rando, 2023, Ma et al., 2015). Progression through the cell cycle presents significant challenges for chromatin organization and epigenetic regulation - especially during S-phase but also during mitosis. Specifically, during S-phase, chromatin must become highly accessible to facilitate replication fork progression, nucleosome redistribution and de novo nucleosome incorporation in the replicated genomes (Stewart-Morgan et al., 2020, Alabert et al., 2015, Reveron-Gomez et al., 2018, Ma et al., 2015, Escobar et al., 2021, Ramos-Alonso and Chymkowitch, 2024). In contrast, in mitosis, chromatin must be tightly compacted to ensure accurate chromosome segregation (Ramos-Alonso and Chymkowitch, 2024, Zhou and Heald, 2020). Central to the regulation of chromatin organization are post-translational histone modifications (PTMs), which are also epigenetic regulators of transcriptional silencing and gene activation (Millan-Zambrano et al., 2022, Bhatnagar et al., 2023, Lopez-Hernandez et al., 2025). It is well-established that PTM dysregulation can lead to cell cycle defects by disrupting chromatin organization and entire gene regulatory networks (Brookes and Shi, 2014, Cavalli and Heard, 2019, Feinberg et al., 2016, Peterson and Laniel, 2004). While much attention has been dedicated to understanding how PTM dysregulation can drive pathogenesis by altering the expression of genes central to cell cycle regulation, less is known about how cell cycle dynamics connect to the maintenance and re-establishment of histone modifications. Yet, studies of the cell cycle-dependent regulation of the histone methyltransferase PR-Set7 and histone demethylase PHF8 underscore the capacity of the cell cycle machinery to control specific histone modifications and thereby maintain chromatin function (Oda et al., 2010, Liu et al., 2010).

Some of the most abundant PTMs that regulate chromatin accessibility and transcriptional activity are acetylation and methylation of lysine residues on histone tails. While several lysine residues in histones can be acetylated or methylated, research has most strongly focused on dynamic acetylation and methylation of lysines 9 and 27 on histone H3 (H3K9 and H3K27), which represent some of the functionally most important histone modifications. In most species, trimethylation of H3K27 is mediated by Polycomb Repressive Complexes and relies on histone methyltransferases of the Enhancer of zeste E(z) family. H3K27me3 maintains the silencing of many genes required for cell fate specification during development, most famously Hox genes (Blackledge and Klose, 2021, Sparmann and van Lohuizen, 2006, Muller et al., 1995, Beuchle et al., 2001, Cao et al., 2002). In contrast, trimethylation of H3K9 (H3K9me3) is mediated by the enzymatic activity of Su(var)3-9 and is central to the formation and maintenance of constitutive heterochromatin at telomeres, centromeres and repetitive repeats (Padeken et al., 2022, Aagaard et al., 1999, Ebert et al., 2004). In *Drosophila*, H3K27 acetylation is mediated by the histone acetyltransferase (HAT) Nejire (Nej), a homolog of mammalian p300/CBP, and is reversed by the histone deacetylase HDAC1/Rpd3 (Willnow and Teleman, 2024, Petruk et al., 2001, Tie et al., 2009, Ciabrelli et al., 2023). H3K27ac is strongly associated with active promoters and enhancers, where it is thought to facilitate chromatin opening and transcriptional activation (Holmqvist and Mannervik, 2013, Philip et al., 2015, Boija et al., 2017, McConnell et al., 2012, Chen et al., 2022). As a consequence of these important functions, dysregulation of H3K9me3, H3K27me3 and H3K27ac is a common feature in cancer or other diseases characterized by alterations and defects in cell proliferation. Yet, importantly, it often remains unclear if defects in epigenetic modifications cause altered proliferation defects or if vice versa proliferation defects cause defects in epigenetic modifications (Brookes and Shi, 2014, Cavalli and Heard, 2019, Feinberg et al., 2016, Classen et al., 2009). Disentangling this relationship is key to understanding how epigenetic and proliferative states influence one another in development and disease.

Importantly, the semi-conservative nature of DNA replication during S-phase has a profound impact on the landscape of these histone modifications. As the DNA is duplicated, pre-existing histones bearing PTMs are equally distributed onto the newly synthesized strands. Naïve, newly synthesized histones are integrated to ensure proper packaging of the duplicated genomes but also necessitating the reestablishment of the now diluted PTM code (Escobar et al., 2021, Alabert and Groth, 2012, Escobar et al., 2019, Wenger et al., 2023). To reestablish the epigenetic histone modification landscape after incorporation of newly synthesized, naive histones, histone-modifying enzymes act post-replication by using the modifications on old recycled histones as a template to modify the newly incorporated histones (Stewart-Morgan et al., 2020, Annunziato, 2015, Flury and Groth, 2024). Importantly, in cultured cells, histone acetylation marks are typically restored immediately after replication often via a transcription-dependent process, while histone methylation is generally re-established during subsequent gap phases (Alabert et al., 2015, Zee et al., 2012, Reveron-Gomez et al., 2018, Petruk et al., 2013).

Of note, strong acetylation of newly synthesized histones plays a specific cell-cycle-dependent role during S-phase. Hyperacetylation, for example at H4K5, H4K12, H3K14, H3K23 or H3K56, is observed during S-phase and facilitates recognition of newly synthesized histones by histone chaperone complexes, promoting their correct incorporation into newly synthesized chromatin. Hyperacetylation of histones presumably occurs in the cytoplasm and may be mediated by B-type acetyltransferases like Hat1 (Li et al., 2008, Masumoto et al., 2005, Tessarz and Kouzarides, 2014, Benson et al., 2006, Falbo and Shen, 2009, Jackson et al., 1976, Lande-Diner et al., 2009, Ruiz-Carrillo et al., 1975, Annunziato and Hansen, 2000, Sobel et al., 1995, Verreault et al., 1998).

Despite these insights, questions remain about how cell cycle progression influences histone modification dynamics, especially in *in vivo* settings. For instance, to what extent are histone modifications regulated in a cell cycle-dependent manner in developing tissues? How does the maintenance of these modifications differ between rapidly proliferating cells and cells in quiescent, senescent, or post-mitotic states? Moreover, how is the activity of histone-modifying enzymes coordinated with cell cycle checkpoints? To begin to address these questions we used the developing *Drosophila* wing imaginal disc to characterize cell cycle dependent histone modifications and reveal pronounced dynamics of H3K9 and H3K27 modifications in a developing tissue.

## RESULTS

### (1) Post-translational modifications of H3K27 and H3K9 are linked to cell cycle dynamics

To understand how cell cycle dynamics may affect chromatin dynamics, we aimed to identify post-translational histone modifications that are sensitive to changes in cell cycle progression. Using the *rn-GAL4* driver **(Fig S1.1A)**, under the control of a temperature-sensitive GAL80ts, we genetically manipulated cell cycle progression in the pouch of developing wing imaginal discs for 24 hours by accelerating or decelerating cell cycle dynamics. Within this timeframe of 24 h, we expect to disturb at least one cell cycle, which lasts between 12 and 18 h in third instar wing imaginal discs (Neufeld et al., 1998, Milan et al., 1996, Prober and Edgar, 2000, Crucianelli et al., 2022). First, we induced a cell cycle arrest in late G2 by expressing *Cdk1-RNAi (Cosolo et al., 2019)*. Second, ectopic expression of the ERK-responsive ETS transcription factor Pointed-P1 (PntP1) causes cells to arrest in either G1 or in G2 (Ito and Igaki, 2021). We confirmed the arrest in our genotypes by observing the cell cycle reporter FUCCI and the absence of EdU incorporation, a nucleotide analogue visualizing DNA replication, in nuclei of the *rn-GAL4* expression domain **(Fig S1.1B-L)** (Zielke et al., 2014). Conversely, we accelerated the cell cycle by co-expressing of dE2F1 and dDP (Buttitta et al., 2007, Neufeld et al., 1998). This did not alter the proportion of cells in G1 or G2 but resulted in elevated EdU incorporation, reflecting the accelerated S-phase dynamics - both in frequency and speed (Crucianelli et al., 2022) - of this genotype **(Fig S1.1M-Q)**.

As post-translational histone modifications are numerous, we focused on prominent modifications mediated by well-known histone-modifying complexes. We analyzed modifications associated with gene regulation, such as H3K27ac (mediated by the p300/CBP nejire), H3K9ac (mediated by SAGA-HAT complex), H3K4me3 (mediated by the Trithorax (Trx)) and H3K27me3 (mediated by the Polycomb protein E(z)), or modifications associated with genome maintenance, such as H3K9me3 (mediated by Su(var)3-9), as well as examples of less well characterized modifications, such as H3K18ac and H4K8ac (mediated by p300/CBP, Gcn5, SAGA-HAT), H3K18 crotonylation (mediated by the p300/CBP nejire), and pan-H3K9 methylation or total lysine acetylation. We performed immunofluorescence staining for these modifications in wing imaginal discs with the induced cell cycle alterations described above.

Many of the histone modifications analyzed did not change upon manipulating cell cycle dynamics. For example, H3K18ac and H3K4me3 levels were not sensitive to cell cycle modulation (**Fig 1A-D, I-L, U-W, Fig S1.3A)**, and neither were H4K8ac, H3K9ac, H3K18 crotonylation, total H3K9 methylation nor levels of total lysine acetylation **(Fig S1.2, Fig S1.3B)**. Yet, strikingly, we found that H3K27ac levels were strongly reduced in cell cycle arrested genotypes, whereas levels of this modification were elevated when the cell cycle was accelerated (**Fig 1E-H, U-W)**. In contrast, H3K9me3 and H3K27me3 levels were elevated in arrested cells, whereas levels of both modifications were reduced when the cell cycle was accelerated (**Fig 1M-W)**. Importantly, alterations observed in cell cycle arrested cells were independent of which cell cycle phase the arrest occurred in. Specifically, H3K27ac reduction and H3K9me3 and H3K27me3 elevation was observed in both G1 and G2 arrested cell populations upon expression of *pntP1* **(Fig S1.1I, Fig 1G,O,S)**, suggesting that the elongation of either gap phase can produce the observed effects.

**Figure 1.**
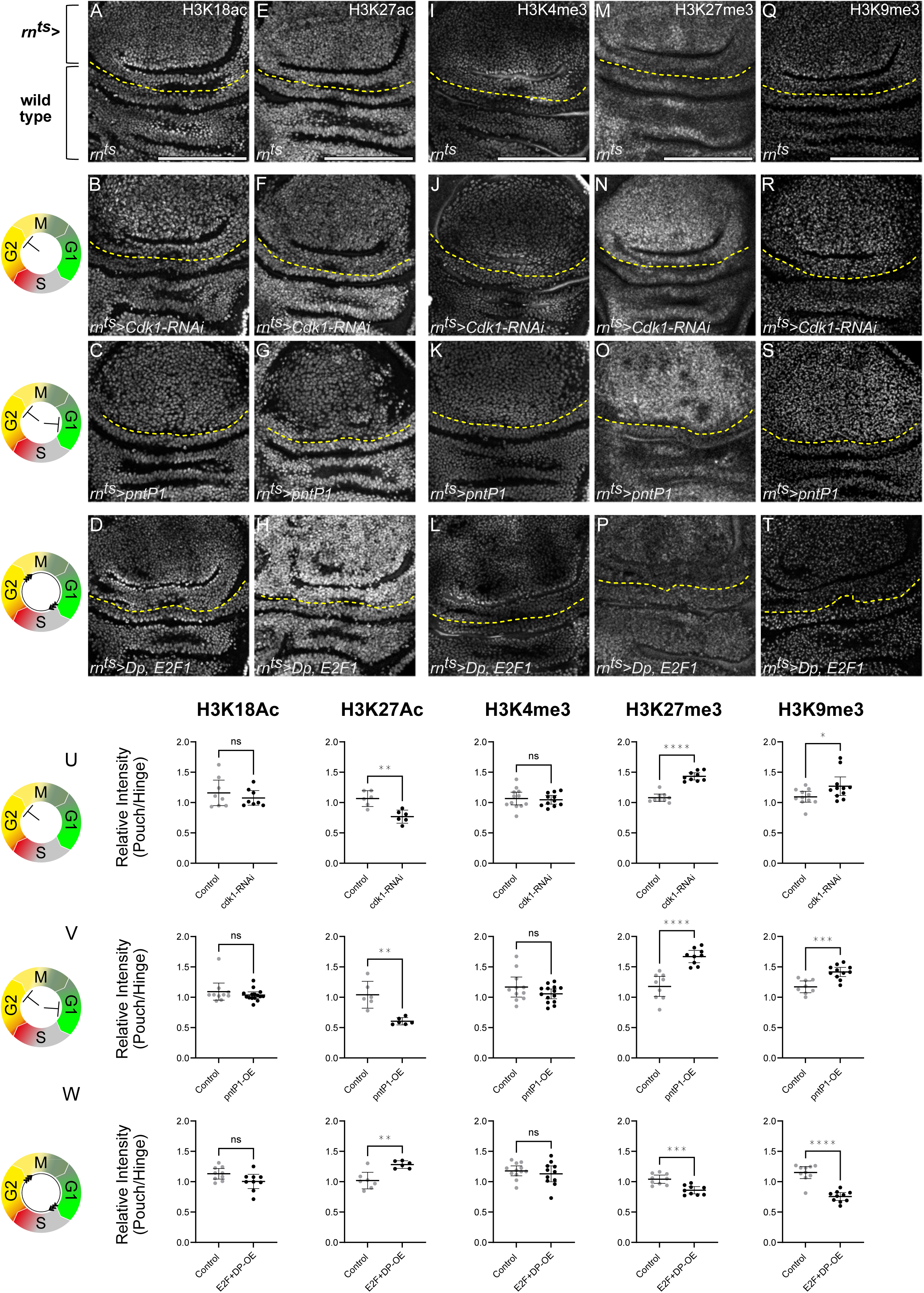
H3K27ac, H3K27me3, and H3K9me3 levels respond to changes in cell cycle dynamics Icons represent the experimentally verified FUCCI cell cycle status in each condition: G2-phase arrest in *Cdk1-RNAi*-expressing discs, either G1 or G2-phase arrest in *pntP1*-expressing discs, and cell cycle acceleration in *Dp, E2F1*-overexpressing discs. For better guidance through subsequent figures, schemes already introduce FUCCI reporter read-outs for phase G1 (green), early S-phase (grey), late S-phase (red) and G2 (yellow). **A-D.** Immunostaining for H3K18ac in control **(A)**, *Cdk1-RNAi*-expressing **(B)**, *pntP1*-expressing **(C)** and *Dp, E2F1*-expressing **(D)** discs. Expression was induced for 24 h in the wing pouch using *rn-GAL4*. The *rn-GAL4* expressing pouch domain is represented by the tissue above the yellow line (rn^ts^>). The tissue below the yellow line represents wild type tissues of the hinge. **E-T**. Immunostaining for H3K27ac **(E-H)**, H3K4me3 **(I-L)**, H3K27me3 **(M-P)**, H3K9me3 **(Q-T)** in control **(E,I,M,Q)**, *Cdk1-RNAi*-expressing **(F,J,N,R)**, *pntP1*-expressing **(G,K,O,S)** and *Dp, E2F1*-expressing **(H,L,P,T)** discs. **U.** Quantification of relative signal intensities for H3K18ac, H3K27ac, H3K4me3, H3K27me3, and H3K9me3, presented as pouch-to-hinge ratios (i.e., rn^ts^-to-wild-type-cell ratio) in *Cdk1-RNAi*-expressing discs. Mean and 95% CI is shown. Statistical significance was tested using two-tailed Mann-Whitney test, p value= 0.9591 for H3K18ac (control discs: n=8, *Cdk1-RNAi*-expressing discs: n=8); two-tailed Unpaired t test, p value=0.0012 for H3K27ac (control discs: n=6, *Cdk1-RNAi*-expressing discs: n=6); two-tailed Unpaired t test, p value=0.7334 for H3K4me3 (control discs: n=12, *Cdk1-RNAi*-expressing discs: n=11); two-tailed Unpaired t test, p value<0.0001 for H3K27me3 (control discs: n=9, *Cdk1-RNAi*-expressing discs: n=9); two-tailed Mann-Whitney test, p value=0.0473 for H3K9me3 (control discs: n=11, *Cdk1-RNAi*-expressing discs: n=11). **V.** Quantification of relative signal intensities for H3K18ac, H3K27ac, H3K4me3, H3K27me3, and H3K9me3, presented as pouch-to-hinge ratios (i.e., rn^ts^-to-wild-type-cell ratio) in *pntP1*-expressing discs. Mean and 95% CI is shown. Statistical significance was tested using two-tailed Mann-Whitney test, p value= 0.5671 for H3K18ac (control discs: n=10, *pntP1*-expressing discs: n=15); two-tailed Welch’s t test, p value=0.0031 for H3K27ac (control discs: n=6, *pntP1*-expressing discs: n=6); two-tailed Unpaired t test, p value=0.1614 for H3K4me3 (control discs: n=11, *pntP1*-expressing discs: n=14); two-tailed Unpaired t test, p value<0.0001 for H3K27me3 (control discs: n=9, *pntP1*-expressing discs: n=9); two-tailed Unpaired t test, p value=0.0002 for H3K9me3 (control discs: n=8, *pntP1*-expressing discs: n=11). **W.** Quantification of relative signal intensities for H3K18ac, H3K27ac, H3K4me3, H3K27me3, and H3K9me3, presented as pouch-to-hinge ratios (i.e., rn^ts^-to-wild-type-cell ratio) in *Dp, E2F1*-coexpressing discs. Mean and 95% CI is shown. Statistical significance was tested using two-tailed Unpaired t test, p value= 0.0609 for H3K18ac (control discs: n=9, *Dp, E2F1*-coexpressing discs: n=9); two-tailed Unpaired t test, p value=0.0019 for H3K27ac (control discs: n=7, *Dp, E2F1*-coexpressing discs: n=6); two-tailed Unpaired t test, p value=0.4837 for H3K4me3 (control discs: n=12, *Dp, E2F1*-coexpressing discs: n=11); two-tailed Unpaired t test, p value=0.0003 for H3K27me3 (control discs: n=9, *Dp, E2F1*-coexpressing discs: n=9); two-tailed Mann-Whitney test, p value<0.0001 for H3K9me3 (control discs: n=10, *Dp, E2F1*-coexpressing discs: n=10). Fluorescence intensities are reported as arbitrary units. Sum projections of multiple confocal sections are shown in **C**, **K**, **M-N**, **O** and **Q-R.** Maximum projections of multiple confocal sections are shown in **A-B**, **E-F**, **G**, **H**, **I-J**, **P**, **S** and **T**. Scale bars: 100 μm. Please refer to **(Fig S1.3A)** for corresponding DAPI images of all discs shown.

Taken together, we conclude that cell cycle dynamics have a surprising impact on histone modification in a proliferating tissue. First, only three tested modifications responded significantly to cell cycle dynamics in our assays, suggesting that only a subset of modifications is strongly linked to the cell cycle machinery. Second, these changes were not dependent to cells arresting in G1 or in G2, suggesting that overall cell cycle length is a defining factor rather than the gap phase position within a cell cycle. Third, if levels of histone modifications were primarily determined by the kinetics of restoring histone modifications after semi-conservative replication and the resulting histone PTM dilution, one may predict that modifications simply accumulate as the cell cycle lengthens. However, the reduction of H3K27ac in arrested cells defies a scenario in which continuous HAT activity restores H3K27ac levels after DNA replication. We thus wanted to better understand the relationship between histone methylation, histone acetylation and cell cycle dynamics.

### (2) Cell cycle dependent changes in H3K27ac, H3K27me3 or H3K9me3 levels occur at preexisting target loci

To first identify the genomic regions affected by the cell-cycle-dependent changes in bulk levels of H3K27ac, H3K27me3, and H3K9me3, we performed CUT&Tag analysis on wing imaginal discs expressing *Cdk1-RNAi* under the control of *rn-GAL4* for 24 hours. Although *Cdk1-RNAi* is expressed in only a subset of cells within the wing disc (see **Fig S1.1A**) - limiting the magnitude of detectable differences - we consistently observed a decrease in H3K27ac levels, and an increase in levels of both H3K27me3 and H3K9me3 modifications. These changes occurred at genomic regions already carrying these modifications in wild type conditions, as evident from examples of genomic snapshots (**Fig 2 A,B)** and changes in average levels of modification on and around peaks identified in both wild type and *Cdk1-RNAi*-expressing discs (**Fig 2 C-F, Fig S2.1A, E)**. These findings are consistent with prior observations in mESCs, where H3K27me3 levels increase at pre-existing sites during prolonged G1 phases (Trouth et al., 2025).

**Figure 2.**
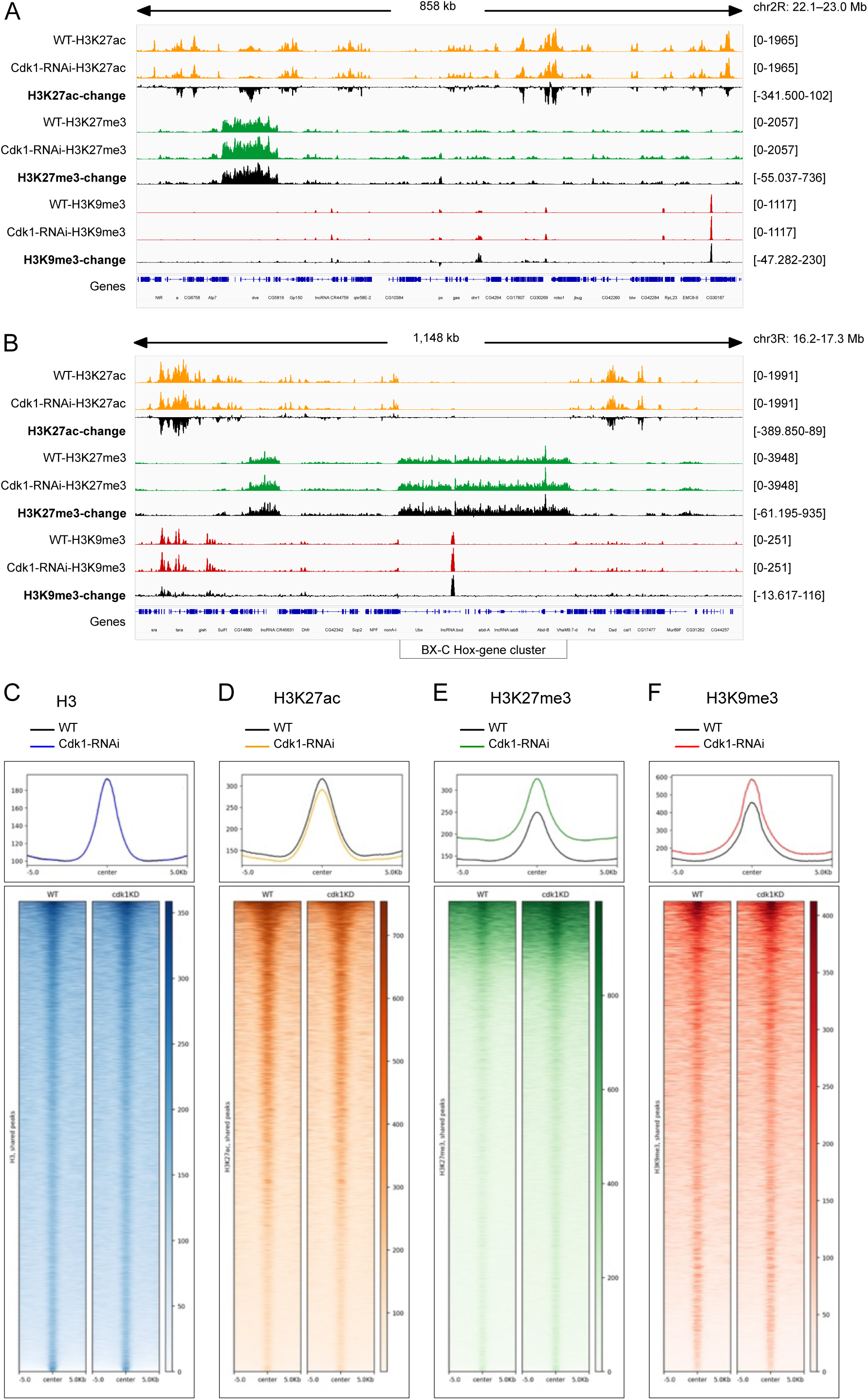
Cell cycle-dependent changes in H3K27ac, H3K27me3, and H3K9me3 occur at preexisting genomic sites **A-B.** Two examples of genomic snapshots of CUT&Tag profiles comparing levels of H3K27ac, H3K27me3 and H3K9me3 in wild type wing discs and in wing discs having expressed *Cdk1*-RNAi for 24 h in the pouch region under the control of rn-GAL4. Tracks for H3K27ac (orange), H3K27me3 (green), and H3K9me3 (red) were normalized against H3 as described in the Materials and Methods and were then visualized using IGV. Black tracks represent the change between control and knockdown conditions computed by subtracting number of sequence reads in *Cdk1-RNAi-*expressing discs from wild type discs for each histone modification. Tracks in **(B)** cover the BX-C Hox gene cluster encoding Ubx, Abd-A and Abd-B targeted by Polycomb silencing. **C-F.** Profile plots and heatmaps showing normalized CUT&Tag signals at peaks shared between wild type control and *Cdk1-RNAi*-expressing discs for H3 **(C)**, H3K27ac **(D)**, H3K27me3 **(E)**, and H3K9me3 **(F)** (see Materials and Methods). Averaged and individual signal intensities are centered ±5 kb around the shared peak.

Of note, many H3K27ac peaks identified only in wild type samples exhibited a significant reduction in H3K27ac levels in *Cdk1-RNAi*-expressing discs, suggesting that H3K27ac was indeed lost from pre-existing acetylated regions **(Fig S2.1B,E)**. In agreement with this conclusion, genomic sequences not identified as peaks by our algorithm in any sample showed little or no change in H3K27ac levels **(Fig S2.1D, E)**. In contrast, although increases in H3K27me3 and H3K9me3 levels in *Cdk1-RNAi* discs occurred at many pre-existing methylated sites **(Fig S2.1A, E)**, a substantial number of H3K27me3 and H3K9me3 peaks identified only in *Cdk1-RNAi* discs, as well as sequences in other genomic regions, also showed significantly elevated methylation compared to wild-type controls **(Fig S2.1C,D,E)**. This indicates that the accumulation of H3K27me3 and H3K9me3 in *Cdk1-RNAi* discs may extend beyond canonical methylation target sites identified in wild type discs. We conclude that cell-cycle-dependent changes to bulk levels of H3K27ac, H3K27me3 or H3K9me3 predominantly affect existing target sites; however, the increase in methylation may also affect other genomic regions. Importantly, the magnitude of changes compared to levels of methylation in control discs is similar for all genomic regions, suggesting that changes to H3K27me3 or H3K9me3 levels occur at the same rate across the genome.

Our conclusions are further supported by our observations that the histone-modifying enzymes responsible for H3K27 acetylation and H3K27methylation do not compete for the same histone residues *in vivo*. Specifically, when we performed a knock-down of the H3K27 methyltransferase E(z), we observed an expected loss of H3K27me3 **(Fig S2.2 A,B)**, but importantly, no corresponding increase in H3K27ac **(Fig S2.2 C,D)**. Conversely, when we performed a knock-down of the H3K27 acetyltransferase CBP/Nej (Nejire), we observed an expected loss of H3K27ac **(Fig S2.2 E,F)**, but no corresponding increase in H3K27me3 **(Fig S2.2 G,H)**. We conclude that, even though studies propose local antagonism of H3K27me3 and H3K27ac at the same residues (Tie et al., 2016, Philip et al., 2015, Tie et al., 2009), the cell-cycle-length-dependent decrease of H3K27ac and increase of H3K27me3 bulk levels upon expression of *Cdk1-RNAi* are not driven by competition for the same H3K27 residues but instead reflect their natural occurrence of these distinct PTMs in distinct genomic contexts.

### (3) H3K27 or H3K9 modifications are linked to cell cycle dynamics during development

If bulk histone modifications in wing discs are affected by cell cycle progression, can we provide evidence that this also occurs in a physiologically relevant state? During larval development, cells in the wing imaginal disc actively proliferate. However, the future wing margin forms a zone of non-proliferating cells (ZNC), where cells arrest in either G1 or G2 **(Fig S3.1 A-C)** (Johnston and Edgar, 1998). In addition to the ZNC, two clusters of G2-arrested cells with senescent markers can be found surrounding the wing pouch **(Fig S3.1 D)** (Zang et al., 2025). Strikingly, arrested cells in the ZNC (**Fig 3 A-C)** and senescent clusters (**Fig 3 D-F)** show a decrease in H3K27ac levels and an increase in H3K9me3 and H3K27me3, consistent with our previous conclusions upon cell cycle elongation by expression of *Cdk1-RNAi* or *UAS-pntP1.* Importantly, none of the other histone modifications recapitulated this pattern, again reinforcing the specific association between H3K9me3, H3K27me3 and H3K27ac levels and the dynamics of cell cycle progression **(Fig S3.1 E-H)**.

**Figure 3.**
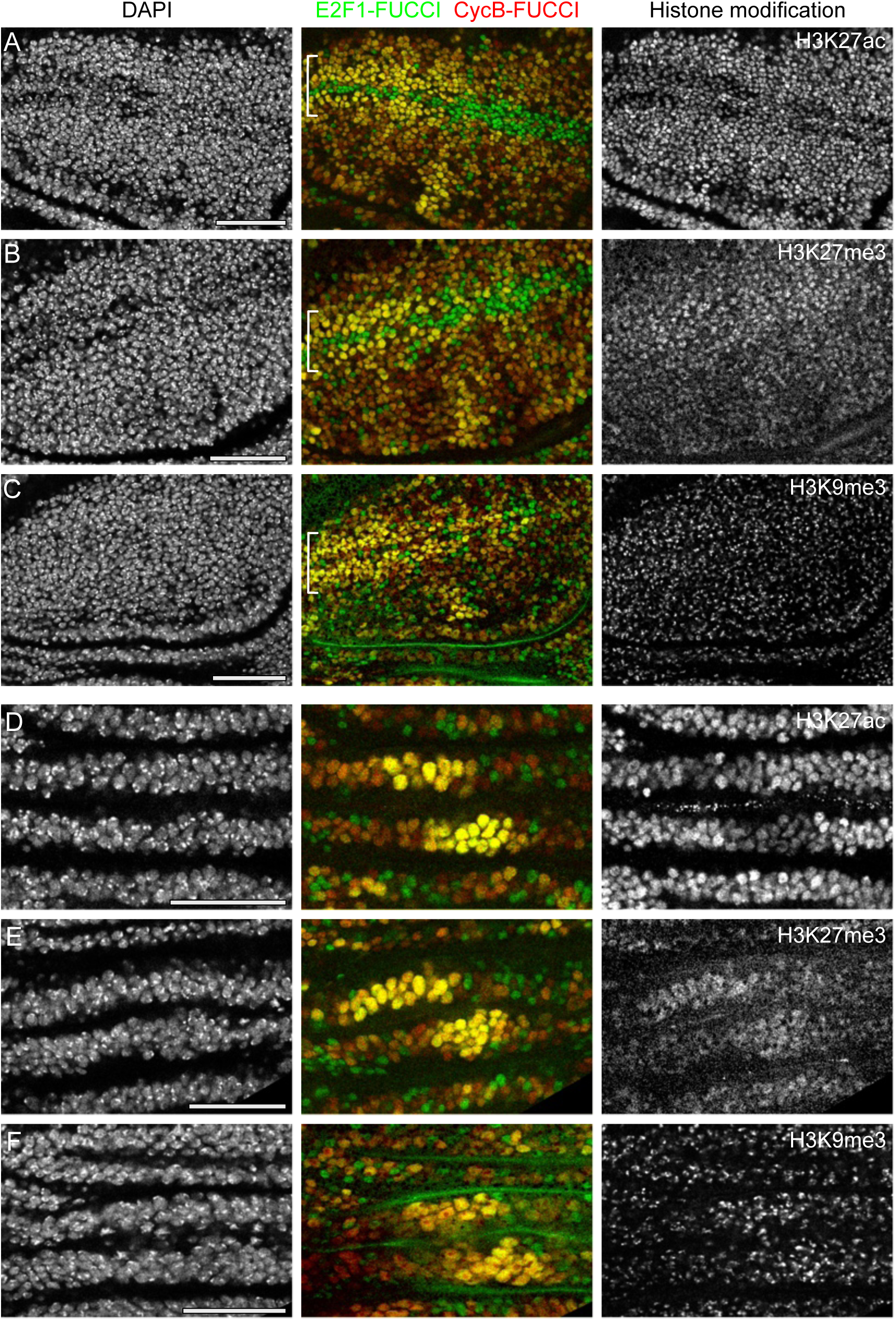
Developmentally arrested cell populations under physiological conditions exhibit changes in H3K27ac, H3K27me3, and H3K9me3 levels **A-C.** Immunostaining for H3K27ac **(A)**, H3K27me3 **(B)**, and H3K9me3 **(C)** in the zone of non-proliferating cells (ZNC) of developing wing discs. Cells in the ZNC (white brackets) are arrested in either G1 or G2 phase, and can be identified by the Fly-FUCCI reporter. Among the histone modifications examined, only H3K27ac, H3K27me3, and H3K9me3 exhibit cell cycle arrest associated changes in the ZNC. **D-F.** Immunostaining for H3K27ac **(D)**, H3K27me3 **(E)** and H3K9me3 **(F)** in a hinge cell population undergoing programmed senescence in developing wing discs. Discs were stained with DAPI to visualize nuclei. Scale bars: 50 μm

To begin to understand if there are consequences for cell-cycle-dependent PTM modulation during development, we analyzed, as an example of three PTMs, H3K27me3-dependent regulation of cell fates. H3K27me3 mediates Polycomb-dependent silencing of genes typically required for cell fate specification (Blackledge and Klose, 2021, Sparmann and van Lohuizen, 2006, Beuchle et al., 2001). In imaginal disc with either arrested (*Cdk1-RNAi*) or accelerated (*Dp,E2F1*) cell cycle dynamics, the expression of Polycomb targeted genes required for wing disc differentiation, such as Antp, Ubx/AbdA, En/inv, Nub, Wg, Ptc and Cut was not altered **(Fig S3.2)**. These observations demonstrate that reduced or elevated H3K27me3 levels observed upon changing cell cycle dynamics still ensure robust Polycomb target gene outputs, suggesting that a H3K27me3 levels buffer gene silencing against a range of cell cycle dynamics.

Surprisingly, our subsequent experiments revealed that the cell cycle-dependent regulation of H3K9me3, H3K27me3, and H3K27ac arises through mechanistically distinct pathways. Specifically, methylation and acetylation marks exhibit fundamentally different dynamics. Given these divergent mechanisms, we present our findings in two consecutive sections: first, we detail how H3K9me3 and H3K27me3 levels are coupled to the cell cycle via growth signaling pathways (**Fig. 4)**, followed by an independent analysis of the function of H3K27ac in preventing replication stress during S-phase (**Figs. 5** and **6)**.

**Figure 4.**
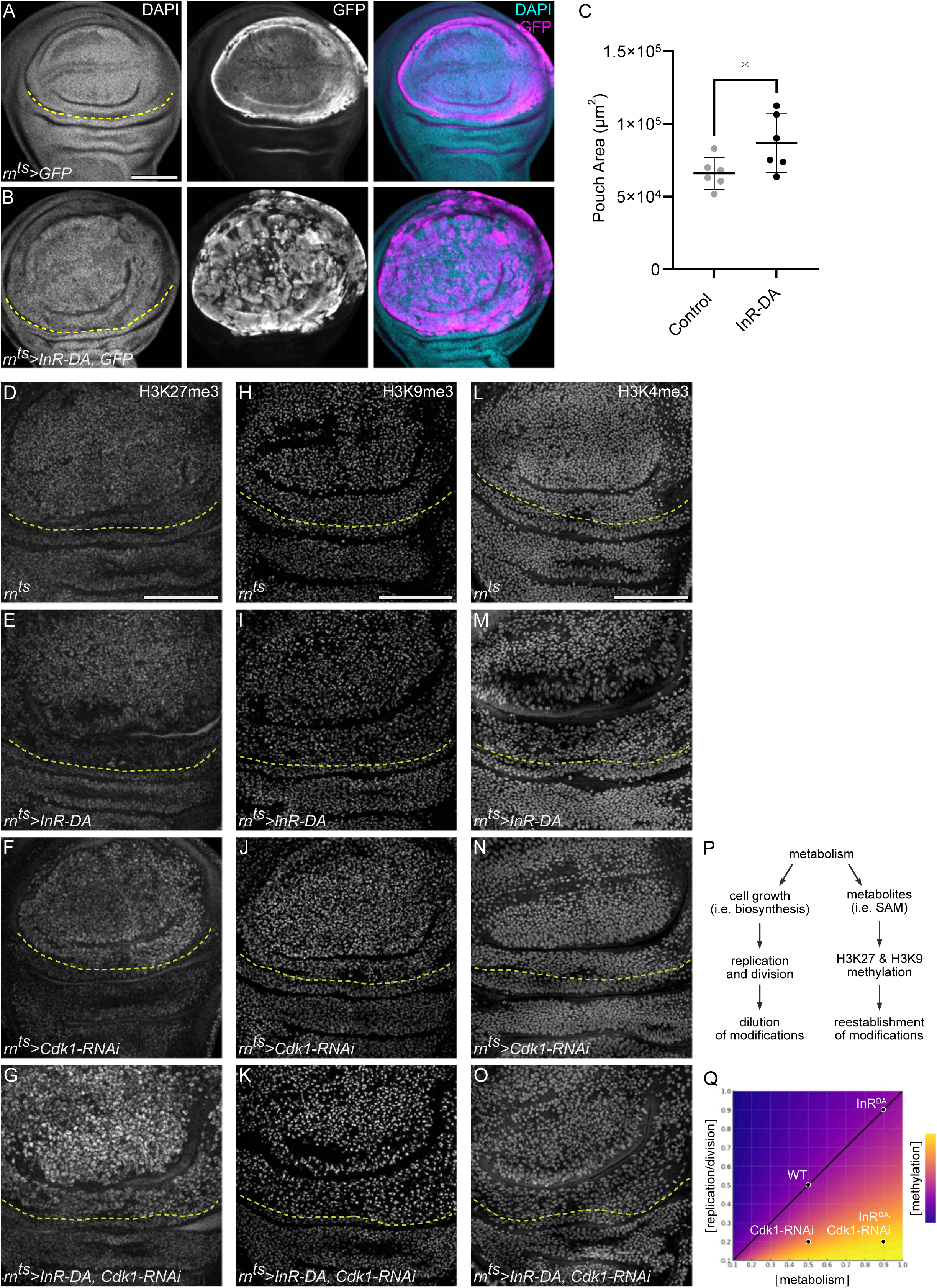
Metabolic state sets the rate of H3K27 and H3K9 methylation **A-B.** Control wing disc expressing GFP (magenta) for 24 h in the wing pouch domain under the control of *rn-GAL4* **(A)**. Wing disc expressing GFP and a constitutively active Insulin receptor (*InR-DA*) under the same conditions **(B)**. GFP expression visualizes the enlarged wing pouch domain in *InR-DA* expressing discs. **C.** Quantification of pouch size area in control and *InR-DA*-expressing discs. Mean and 95% CI is shown. Statistical significance was tested using two-tailed Unpaired t test, p value=0.0432. (control discs: n=6, *InR-DA-*expressing discs: n=6). **D-G.** Immunostaining for H3K27me3 in control **(D)**, *InR-DA*-expressing **(E)**, *Cdk1-RNAi*-expressing **(F)** and *InR-DA, Cdk1-RNAi*-coexpressing **(G)** discs. **H-K.** Immunostaining for H3K9me3 in control **(H)**, *InR-DA*-expressing **(I)**, *Cdk1-RNAi*-expressing **(J)** and *InR-DA, Cdk1-RNAi*-coexpressing **(K)** discs. **L-O.** Immunostaining for H3K4me3 in control **(L)**, *InR-DA*-expressing **(M)**, *Cdk1-RNAi*-expressing **(N)** and *InR-DA, Cdk1-RNAi*-coexpressing **(O)** discs. Expression was induced for 24 h in the wing pouch using *rn-GAL4.* Yellow dashed lines indicate the boundary between the wing pouch and hinge regions. The area above the line corresponds to the *rn-GAL4* expression domain, while the area below represents wild-type cells. **P.** Illustration of the dual role of metabolism in regulating histone methylation levels. Metabolism contributes to the dilution of histone PTMs by accelerating the cell cycle, thereby increasing the frequency of DNA replication events. Simultaneously, it supports the re-establishment of these PTMs by supplying essential precursors and cofactors required for methyltransferase activity. **Q.** Hypothetical model depicting the relationship between histone methylation levels, metabolic rate, and cell division rate. When metabolic activity and cell division are proportionally coupled (black diagonal), histone methylation levels are maintained at stable levels. However, when metabolism outpaces cell division, methylation accumulates. Discs were stained with DAPI to visualize nuclei. Sum projections of multiple confocal sections are shown in **A-B.** Maximum projections of multiple confocal sections are shown in **D-K.** Scale bars: 100 μm.

**Figure 5.**
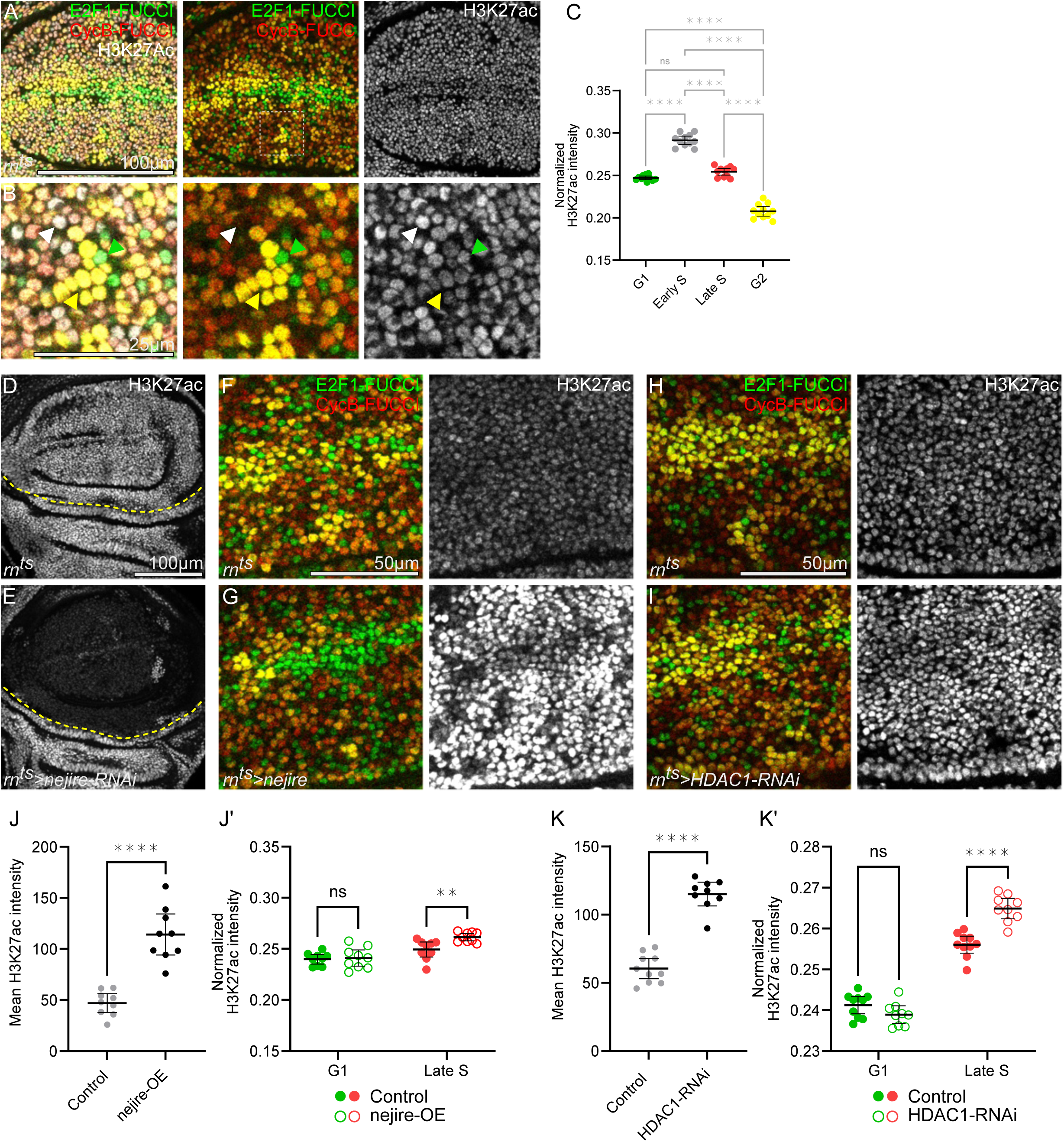
CBP/nejire-mediated H3K27 acetylation peaks during S phase **A-B.** Immunostaining for H3K27ac in a control disc expressing the FUCCI reporters *GFP-E2F1*^1-230^ (green) and *mRFP-NLS-CycB*^1-266^ (red) to visualize cell cycle phases **(A).** Magnified region in **(B)** is demarcated by a white dashed square in **(A). C.** Quantification of normalized H3K27ac intensity level in different cell cycle phases in the pouch region of control discs. Normalization was performed against the average fluorescence intensity across all cell cycle phases. Mean and 95% CI is shown. Statistical significance was tested using Repeated Measures One-Way ANOVA followed by Tukey’s post-hoc test for multiple comparison (n=11 discs). Adjusted p<0.0001 for G1 vs. Early S; p=0.0838 for G1 vs. Late S; p<0.0001 for G1 vs. G2; p<0.0001 for Early S vs. Late S; p<0.0001 for Early S vs G2 and p<0.0001 for Late S vs G2. **D-E.** Immunostaining for H3K27ac in control **(D)** and *nejire-RNAi*-expressing **(E)** discs. Expression was induced for 24 h in the wing pouch using *rn-GAL4.* Yellow dashed lines indicate the boundary between the wing pouch and hinge regions. **F-G.** Immunostaining for H3K27ac in control **(F)** and *nejire*-expressing **(G)** discs also expressing the FUCCI reporters *GFP-E2F1*^1-230^ (green) and *mRFP-NLS-CycB*^1-266^ (red) to visualize cell cycle phases. Expression of *nejire* was induced for 24 h in the wing pouch using *rn-GAL4*. **H-I.** Immunostaining for H3K27ac in control **(H)** and *HDAC1-RNAi*-expressing **(I)** discs also expressing the FUCCI reporters *GFP-E2F1*^1-230^ (green) and *mRFP-NLS-CycB*^1-266^ (red) to visualize cell cycle phases. Expression of *HDAC1-RNAi* was induced for 24 h in the wing pouch using *rn-GAL4*. **J.** Quantification of mean H3K27ac intensity in the pouch region of control or *nejire*-expressing discs. Expression was induced for 24 h in the wing pouch using *rn-GAL4.* Mean and 95% CI is shown. Statistical significance was tested using two-tailed Welch’s t test, p value<0.0001. (control discs: n=9, *nejire-* expressing discs: n=9). **J’.** Quantification of normalized H3K27ac in G1 phase and Late S phase in the pouch region of control or *nejire*-expressing discs. Expression was induced for 24 h in the wing pouch using *rn-GAL4.* Mean and 95% CI is shown. Statistical significance was tested using Repeated Measures Two-Way ANOVA followed by with Šidák’s post hoc test for multiple comparisons (control discs: n=9, *nejire*-expressing discs: n=9). Adjusted p= 0.9482 for Control G1 vs nejire-OE G1; p=0.0064 for Control Late S vs nejire-OE Late S. **K.** Quantification of mean H3K27ac intensity in the pouch region of control or *HDAC1-RNAi*-expressing discs. Expression was induced for 24 h in the wing pouch using *rn-GAL4.* Mean and 95% CI is shown. Statistical significance was tested using two-tailed Mann-Whitney test, p value<0.0001. (control discs: n=10, *HDAC1-RNAi-*expressing discs: n=9). **K’.** Quantification of normalized H3K27ac intensity level in G1 and Late S phases in the pouch region of control or *HDAC1-RNAi*-expressing discs. Expression was induced for 24 h in the wing pouch using *rn-GAL4.* Mean and 95% CI is shown. Statistical significance was tested using Repeated Measures Two-Way ANOVA followed by with Šidák’s post hoc test for multiple comparisons (control discs: n=10, *HDAC1-RNAi*-expressing discs: n=9). Adjusted p= 0.1907 for Control G1 vs HDAC1-RNAi G1; p<0.0001 for Control Late S vs HDAC1-RNAi Late S. Fluorescence intensities are reported as arbitrary units. Sum projections of multiple confocal sections are shown in **D-E.** Scale bars: 100 μm in **A** and **D-E**; 50 μm in **F-G** and **H-I**; 25 μm in **B**.

**Figure 6.**
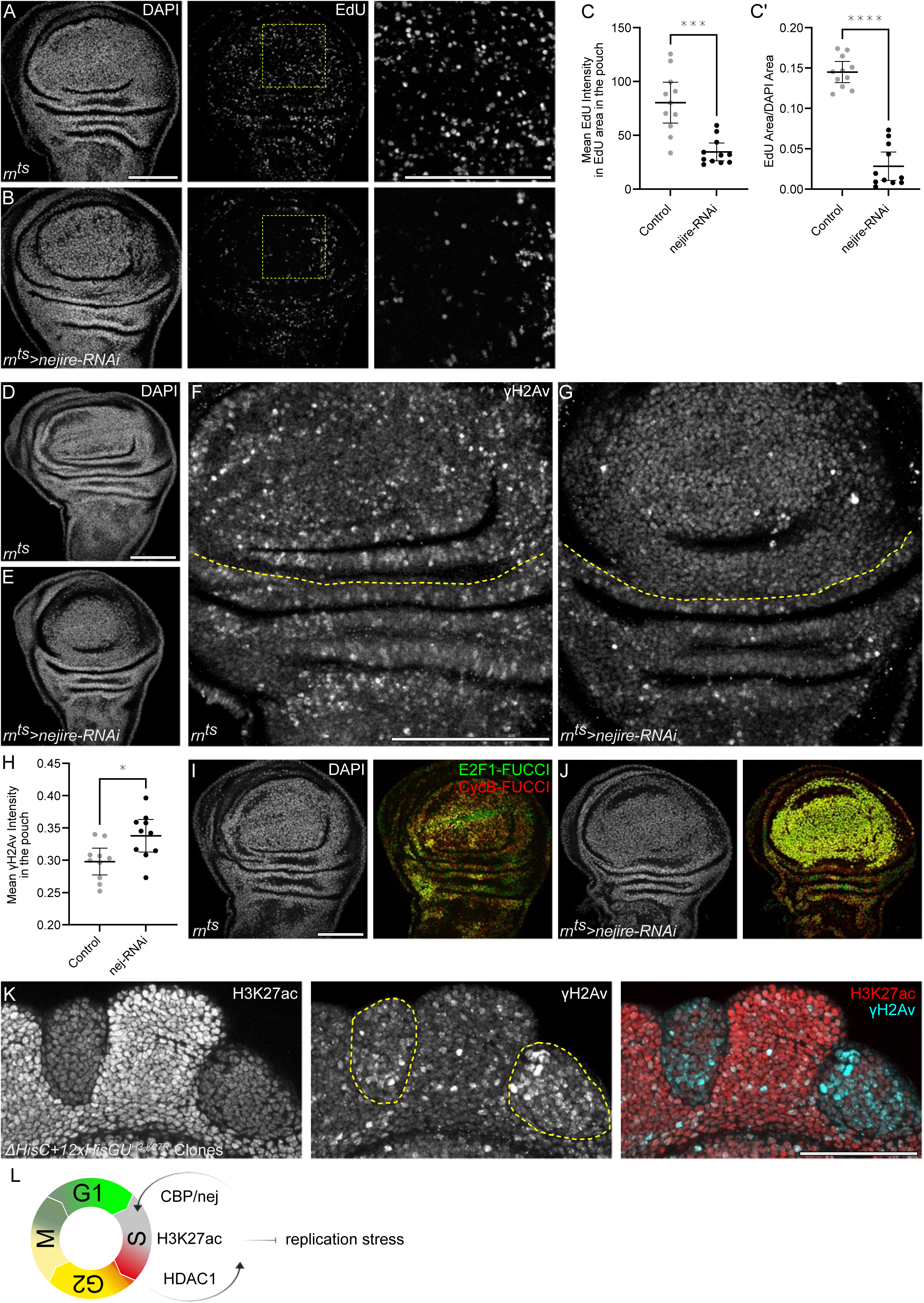
CBP/Nejire activity and H3K27 acetylation prevent replication stress during S phase **A-B.** EdU incorporation to visualize DNA replication in the pouch region of control **(A)** and *nejire-RNAi-* expressing **(B)** discs. Dashed yellow squares demarcate magnified regions shown in **(A’, B’).** Expression was induced for 24 h in the wing pouch using *rn-GAL4*. **C.** Quantification of mean EdU intensity per EdU area in the pouch region of control and *nejire-RNAi*-expressing discs, serving as a proxy speed of DNA replication. Mean and 95% CI are shown. Statistical significance was tested using two-tailed Mann Whitney test, p value=0.0001 (control discs: n=11, *nejire-RNAi*-expressing discs: n=11). **C’.** Quantification of EdU area per DAPI area in the pouch region of control and *nejire-RNAi*-expressing discs, serving as a proxy number of cells undergoing DNA replication. Mean and 95% CI are shown. Statistical significance was tested using two-tailed Mann Whitney test, p value<0.0001 (control discs: n=11, *nejire-RNAi*-expressing discs: n=11). **D-G.** Immunostaining for γH2Av in control **(F)** and *nejire-RNAi*-expressing **(G)** discs. Expression was induced for 24 h in the wing pouch using *rn-GAL4.* γH2Av serves as a marker of DNA double strand breaks. DAPI staining was used to visualize the nuclei in control **(D)** and *nejire-RNAi-*expressing **(E)** discs. The *rn-GAL4* expression domain is represented by the tissue above the yellow line (rn^ts^>). The tissue below the yellow line represents wild type tissues of the hinge. **H.** Quantification of mean γH2Av intensity in the pouch region of control or *nejire-RNAi*-expressing discs. Mean and 95% CI are shown. Statistical significance was tested using two-tailed Unpaired t test, p value=0.0126 (control discs: n=10, *nejire-RNAi*-expressing discs: n=10). **K.** Immunostaining for H3K27ac (red) and γH2Av (cyan) in wing imaginal discs with clones of *ΔHisC* mutant clones deleting the histone cluster and rescued by a *12×His-GU^H3-K27R^* transgene. **L.** Illustration of the role of CBP/nej and HDAC1 in modulating levels of H3K27ac during S-phase to prevent replication stress Discs were stained with DAPI to visualize nuclei. Fluorescence intensities are reported as arbitrary units. Sum projections of multiple confocal sections are shown in **A-B** and **D-G.** Maximum projections of multiple confocal sections are shown in **K.** Scale bars: 100 μm in **A-B**, **D-G**, **I-J** and **K.**

### (4) H3K27me3 or H3K9me3 are coupled to cell cycle dynamics via metabolic signaling

Previous studies suggest that H3K27me3 and H3K9me3 modifications after DNA-replication are reestablished after DNA replication through slow and continuous methyltransferase activity in subsequent gap phases (Alabert et al., 2015, Zee et al., 2012, Reveron-Gomez et al., 2018). Our findings are consistent with these observations: in all our imaginal disc models, levels of H3K9me3 and H3K27me3 increase in conditions with elongated cell cycles (and thus infrequent S-phase events, reducing replication-associated dilution of these marks), and decrease in conditions with shortened cell cycles (increasing the frequency of S-phase and thereby enhancing dilution). These results suggest that H3K9 and H3K27 methylation is driven by a relatively constant rate of histone methyltransferase activity, which appears not to be affected by genetic alterations of the cell cycle machinery. If the rate of histone methylation can indeed be genetically separated from the frequency of cell cycling, what mechanisms then govern the restoration of histone methylation to the appropriate levels, particularly for H3K9me3, which is essential for heterochromatin architecture, and H3K27me3, which is critical for gene silencing? To address this question, we decided to manipulate cell cycle progression upstream of direct regulators of the cell cycle by activating a proliferative pathway with important metabolic functions. Specifically, we expressed a constitutively active Insulin receptor (InR^DA^) under the control of *rn-GAL4* for 24 h, which promotes PI3K/Akt signaling. The resulting metabolic changes can be visualized by higher rates of protein synthesis reflecting higher biosynthetic capacity, accelerated cell proliferation visualized by high EdU incorporation and ultimately, larger wing pouch size (**Fig 4A-C, Fig S4 A-E)**. Of note, expression of InR^DA^ did not alter the FUCCI cell cycle profile, demonstrating that the higher metabolic activity shortens the overall cell cycle length, rather than affecting specific cell cycle phases **(Fig S4F,G)**.

Importantly, we found that expression of InR^DA^ alone did not give rise to changes in H3K27 or H3K9 methylation levels, despite the accelerated cell divisions (**Fig 4 D,E,H, I)**. This demonstrates that the metabolic potential can tune the rate of H3K27 and H3K9 methylation, to keep levels of H3K27 or H3K9 methylation comparable to wild type levels, despite accelerated cell cycle progression. To now functionally separate metabolic potential from cell cycle progression, we also arrested the cell cycle by co-expressing the constitutively active Insulin receptor with a *Cdk1-RNAi*. As expected from our previous experiments, levels of H3K27me3 and H3K9me3 increased, but most strikingly, levels of H3K27me3 and H3K9me3 were even higher than in discs expressing *Cdk-1-RNAi* alone (**Fig 4F,G,J,K)**. These results demonstrate that rates of H3K27 or H3K9 methylation are intimately linked to the metabolic potential of a cell - with the purpose to match the rate of H3K27 or H3K9me3 methylation with the speed of cell cycle progression and thus the frequency of S-phase linked histone dilution (**Fig 4 P,Q)**. Of note, levels of H3K4me3 were not affected by these manipulations (**Fig 4 L-O)**, indicating that a modification maintaining steady levels of bulk transcriptional activity is not - and even should not be - linked to metabolic rates or the rate of cell cycle progression.

We wanted to better understand the functional implications of our findings. The link between levels of H3K9me3 and cell cycle progression becomes relevant in physiological and pathological setting of senescence, where cytotoxic stress or oncogenic mutations provide metabolic activation that is juxtaposed to a senescent cell cycle arrest program. Accordingly, in senescent G2-arrested cells at the center of inflammatory tissue damage in wing imaginal discs induced by ectopic expression of the TNFα homologue eiger (egr) **(Fig S4.2 A-F)** (Cosolo et al., 2019, Floc’hlay et al., 2023, Jaiswal et al., 2023, Smith-Bolton et al., 2009), we observed high levels of H3K9 methylation **(Fig S4.2G-I, purple)**. Importantly, levels in senescent cells with inflammatory metabolism were higher than in quiescent cells at the disc periphery, which exhibit low insulin-signaling activity **(Fig S4.2G-I, yellow)** (Maya et al., 2024). These observations support our conclusions that high and low metabolic activity coupled to cell cycle arrest produce different levels of H3K9me3. We thus propose that the widely reported senescence-associated heterochromatinization associated with elevated H3K9me3 during oncogene-induced senescence may not be a feature of the senescence program *per se* but rather a consequence of continuous H3K9 trimethylation driven by metabolic activation from oncogenic mutations in the absence of cell division (Hernandez-Segura et al., 2018, Huang et al., 2022, Narita et al., 2003). Finally, in agreement with our previous conclusions, we found that cells undergoing fast regenerative proliferation around the site of inflammatory damage have lowest levels of H3K9me3 **(Fig S4.2G-I, cyan)**.

### (5) CBP/nejire-mediated H3K27 acetylation is high during S-phase

Having revealed how H3K9me3 and H3K27me3 dynamics are coupled via metabolic cues to the cell cycle, we now wanted to characterize the observed cell cycle dynamics of H3K27 acetylation. The downregulation of H3K27ac by cell cycle elongation and the upregulation of H3K27ac by cell cycle acceleration contradicts the simple explanation that an enzymatic activity acetylates H3K27 at a constant rate. To better understand H3K27ac dynamics, we performed a detailed analysis of H3K27ac levels throughout the cell cycle by correlating H3K27ac with FUCCI reporters in wing imaginal discs. We specifically excluded mitotic cells from this analysis, as the highly condensed nature of their chromosomes and the cells’ extreme apical position prevented a direct comparison to other cell cycle stages.

Using this approach, we found that H3K27ac levels were significantly increased in nuclei of early and late S-phase cells if compared to G1 and G2 phases (**Fig 5A-C)**. This increase, and especially the much higher levels in late S-phase in comparison to G2, could only partially be recapitulated by changes in total H3 or total acetyl-Lysine levels **(Fig S5.1 A-F)**. Of note, fluctuation of H3 and total lysine acetylation in our microscopy assays can be mostly explained by S-phase dependent histone synthesis and histone acetylation, as well as fluctuating morphology of interphase nuclei (Stein et al., 1975, Mendiratta et al., 2019, Verreault et al., 1998). Our finding that H3K27ac levels are higher throughout S-phase are consistent with our earlier observations: cells which cycle fast repeatedly enter S-phase and cells which cycle slowly are not often found in S-phase, hence bulk H3K27ac levels in a proliferating and arrested tissue will appear high and low, respectively.

These results suggest that the enzymes responsible to regulate H3K27 acetylation may exhibit cell-cycle-dependent regulation. Knock-down of the H3K27ac HAT CBP/nej via RNAi (*neijire-RNAi*) expressed for 24 h under the control *rn-GAL4*, completely abolished H3K27ac staining in the wing pouch, confirming that CBP/nej is the main source of H3K27 acetylation in the wing disc (**Fig 5 D-E)** (Willnow and Teleman, 2024). To provide specific proof that CBP/nej mediates S-phase-dependent H3K27 acetylation, we overexpressed CBP/nej for 24 h under the control of *rn-GAL4* and correlated H3K27ac levels with cell-cycle phases using the FUCCI reporter. While global H3K27ac increased upon CBP/nej overexpression (**Fig 5F,G,J; Fig S5.1 G,H)**, we observed a disproportionate increase of H3K27ac in late S-phase cells (**Fig 5J’)**. This demonstrates that CBP/nej-driven acetylation of H3K27 is especially pronounced during S-phase and indicates that H3K27ac accumulates ectopically in late S-phase cells when CBP/nej levels are too high. In support of a model where CBP/nej is specifically active during S-phase, we found that levels of two other CBP/nej-mediated modification (H3K18ac and H4K8ac) showed a less pronounced but still detectable increase in S-phase cells of normally proliferating wild type tissue **(Fig S5.2 A-F)**, which is consistent with the reduced CBP/nej substrate specificity for these two residues, as well as for H3K18 crotonylation, observed in CBP/nej-RNAi experiments **(Fig S5.2 G-N)**.

As these targeted overexpression experiments suggested that CBP/nej levels may influence S-phase-dependent activity, we examined a CBP/nej-GFP fusion protein expressed from the endogenous locus to assess if CBP levels fluctuate during the cell cycle (Marsh et al., 2025). We found that, in proliferating wing discs, nuclear GFP levels were mostly uniform in all cells **(Fig S5.2 O)**, indicating that regulation of CBP/nej enzymatic activity, rather than cell cycle dependent changes in protein levels, underlies the S-phase-specific activity. Indeed, in cultured cells, CBP/p300 activity may be regulated by cyclin-dependent kinases at the G1/S transition (Ait-Si-Ali et al., 1998).

The pronounced reduction of H3K27ac in late-S-phase, and subsequently in G2-phase or G1-phase cells, which we observed in development, in arrested cell populations and upon CBP/nej overexpression or knock-down, demonstrates that the removal of H3K27ac in late S-phase must be a highly controlled process. Removal of acetylation from the H3K27 residue would likely be mediated by HDAC activity. We thus depleted the primary H3K27 deacetylase HDAC1 (Rpd3) via expression of an RNAi construct under the control of *rn-GAL4* for 24 h (Willnow and Teleman, 2024). HDAC1 knockdown led to a mild but detectable overall increase in H3K27ac, consistent with its role in H3K27ac removal (**Fig 5 H, I, K; Fig S5.1 I,J)**. Yet, importantly, we observed a disproportionate increase of H3K27ac in late S-phase cells, supporting a model where HDAC1 is required to specifically remove H3K27ac in late S-phase (**Fig 5K’)**.

### (6) CBP/nej activity and H3K27acetylation is required to prevent replication stress during S-phase

We finally wanted to understand what function elevated CBP/nej activity during S-phase may serve. To analyze if elevated CBP/nej activity may reflect a specific requirement during DNA replication, we conducted a knock-down of *CBP/nej* in the wing pouch under the control of *rn-GAL4* for 24 h. We first analyzed DNA replication by EdU incorporation and found a wide-spread decrease in the number of cells entering S-phase and in DNA replication activity, suggesting that both S-phase entry and progression was severely slowed within 24 h of *CBP/nej* knock-down (**Fig 6A-C)**. To determine if DNA replication during *CBP/nej* knock-down had proceeded successfully, we investigated if *CBP/nej-RNAi*-expressing cell experienced replication stress. We thus analyzed levels of phosphorylated γH2A, a marker of DNA double strand breaks which can arise at stalled replication forks (Madigan et al., 2002, Petermann et al., 2010). In all wild type tissues, we observed a similar number of high intensity nuclei positive for γH2A, likely representing physiological cell death or (unspecific) binding of the antibody in mitotically active cells (Khan et al., 2017). However, the majority of nuclei are negative for γH2A (**Fig 6D,F)**. Strikingly, in all nuclei of *CBP/nej-RNAi* - expressing cells, we observed a pronounced increase in nuclear γH2A levels - which were represented by medium fluorescence intensities. The γH2A signal covered the entire nuclei, suggesting wide-spread DNA damage (**Fig 6E,G,H)**. We conclude that CBP/nej is specifically required to prevent genome-wide replication stress during S-phase. Accordingly, *CBP/nej-RNAi*-expressing cells cannot progress through the cell cycle and completely arrest in G2 (**Fig 6I,J)**. In tissue damage models of wing discs, cells arrest in G2 as a consequence of stress-dependent activation of JNK-signaling, which downregulates *cdc2/stg* (Cosolo et al., 2019). Yet, co-expressing a dominant-negative JNKK (bskDN) in *CBP/nej-RNAi*-expressing caused pronounced downregulation of JNK activation **(Fig S6A-C)**, but still resulted in G2 arrested cells **(Fig S6D)**. This strongly indicates that a very specific function of CBP/nej in DNA replication caused a G2 cell cycle arrest by activation of a DNA damage checkpoint, rather than the non-specific activation of cellular stress signaling through JNK. These observations are thus consistent with a model were high levels of CBP/nej-mediated H3K27ac are specifically required to ensure faithful DNA replication and to prevent DNA damage and consequently, a DNA damage induced cell cycle arrest in G2. Of note, analysis of *CBP/nej-*overexpressing tissues revealed normal patterns of EdU incorporation and distribution, suggesting that cells with high - but still cell-cycle-dependent - levels of H3K27ac proliferate normally **(Fig S6E-G)**.

CBP/nej targets a wide range of substrates, including several non-histone proteins (Stauffer et al., 2007, Cazzalini et al., 2014, Dutto et al., 2018, Tillhon et al., 2012, Ramadan et al., 2025, Ogiwara et al., 2011). We thus wanted to determine, if the function of CBP/nej in preventing replication stress is mediated by its substrate H3K27. Especially, since H3K27 was the histone substrate most sensitive to the loss of CBP/nej activity in our assays (refer to **Fig S5.2 G-N)**. To test this, we generated mosaic clones homozygous for a deletion of the entire histone gene cluster (ΔHisC), whose viability was rescued by 12 copies of a histone gene unit in which wild type H3 was replaced with a non-modifiable H3K27R amino acid substitution (Pengelly et al., 2013). As previously reported, these clones lose Polycomb gene silencing function normally mediated by H3K27me3, and exhibit derepression of Hox genes such as *AbdB* **(Fig S6H,I)** (Pengelly et al., 2013). When we assessed if these cells also exhibit replication stress, we observed broad and elevated levels of γH2A in the H3K27R-substituted clones (**Fig 6K)**. These findings suggest that the function of CBP/nej in mitigating replication stress during DNA replication may be, at least in part, mediated through its modification of H3K27 (**Fig 6I)**.

## DISCUSSION

Our work dissects the link between H3K27ac, H3K27me3, and H3K9me3 modification dynamics and dynamics of the cell cycle in proliferating tissues *in vivo*. We demonstrate that both physiological and genetically induced elongation of the cell cycle leads to elevated bulk levels of H3K27me3 and H3K9me3. Conversely, accelerated cell cycle progression specifically decreases these histone marks. Genome-wide analyses using CUT&tag assays revealed that these changes occur predominantly at their distinct pre-existing genomic target loci, implicating canonical histone readers and modifiers in mediating these cell cycle-dependent changes. Importantly, despite cell cycle-dependent changes in bulk H3K27me3 levels, expression patterns of Polycomb target genes remained unaffected, implying that H3K27me3-dependent gene silencing at target loci is buffered even under altered cell cycle conditions. Mechanistically, we found a tight coupling between histone methylation rates and cell division frequency, mediated by cellular metabolic activity. Specifically, H3K27me3 and H3K9me3 levels remained normal upon expression of a constitutively active Insulin receptor, suggesting that cells channeled the higher metabolic activity into accelerated cell divisions and maintained H3K27me3 and H3K9me3 levels proportional to cell division rates. Remarkably, active Insulin receptor-expressing cells forced to arrest their cell cycle accumulated levels of H3K27me3 and H3K9me3 to much higher levels than arrested wild type cells, demonstrating that metabolic rate tunes the rate of histone methylation to match the frequency of S-phase occurrence, i.e. the frequency of cell divisions. These observations have important implications for chromatin dysregulation in pathological conditions, such as cancer, where the tight coordination between metabolic and cell cycle rates becomes compromised. An important implication of our findings relates to the emergence senescence-associated heterochromatin, a phenomenon well-documented during oncogene-induced senescence (Hernandez-Segura et al., 2018, Huang et al., 2022, Narita et al., 2003). Our data suggest that H3K9me3 accumulation in senescence-associated heterochromatin may reflect persistent methyltransferase activity driven by aberrant metabolic stimuli in the absence of replication-dependent dilution. Thus, the observed chromatin state in senescent cells may be a consequence of the mismatch between metabolic and cycling rate, rather than an intrinsic feature of the senescence phenotype itself. An important implication of our findings is that post-mitotic and differentiating cells, such as neurons, must tightly regulate histone-modifying activities to prevent the aberrant accumulation of repressive marks, such as H3K9me3 and H3K27me3.

Furthermore, we observed that H3K27ac exhibits a unique, cell cycle–dependent profile, peaking significantly during S-phase. While hyperacetylation of newly synthesized histones at various residues during S-phase is well-established, an S-phase dependent increase in nuclear acetylation of H3K27 has not been reported (Li et al., 2008, Masumoto et al., 2005, Tessarz and Kouzarides, 2014, Benson et al., 2006, Falbo and Shen, 2009, Jackson et al., 1976, Lande-Diner et al., 2009, Ruiz-Carrillo et al., 1975, Annunziato and Hansen, 2000, Sobel et al., 1995). We propose this peak in acetylation during S-phase is mediated by CBP/nej, potentially regulated by cyclin-dependent kinases at the G1/S transition (Ait-Si-Ali et al., 1998). Correspondingly, our data is consistent with the idea that H3K27 becomes deacetylated during late S-phase by HDAC1. Generally, HDAC activity is reported to remove hyperacetylation on newly synthesized histone at the end of S-phase to reestablish higher order chromatin organization (Annunziato and Seale, 1983). However, whether the H3K27 acetylation cycle mediated by CBP and HDAC 1 occurs on old histones, newly synthesized histones, or on newly incorporated histones cannot be resolved with our methods. Yet, the observed H3K27 acetylation cycle reveals novel cell cycle dependent targets of CBP/nej and HDAC1.

Functionally, we demonstrate that CBP/nej activity is essential during S-phase to prevent DNA damage from replication stress and consequently delays in cell cycle progression mediated by a replication checkpoint. In fact, previous studies find that CBP/nej and ATR mei-41 are required for the replication checkpoint itself to function properly (Smolik and Jones, 2007, Stauffer et al., 2007). This may be explained by acetylation non-histone substrates by CBP/p300, which stabilizes key DNA damage repair factors, including PCNA, ATR and ATM, thereby facilitating DNA repair following damage (Stauffer et al., 2007, Cazzalini et al., 2014, Dutto et al., 2018, Tillhon et al., 2012, Ramadan et al., 2025, Ogiwara et al., 2011). Consistent with this, mutations in EP300 result in persistent replication stress, replication fork collapse, genomic instability, and embryonic proliferation defects (Barreto-Galvez et al., 2023; Yao et al., 1998). (Barreto-Galvez et al., 2023, Yao et al., 1998). Collectively, these findings underscore the essential role of CBP/p300-mediated acetylation in regulating replication integrity and genomic stability beyond traditional cell cycle regulation. Importantly, our findings extend this previous work by providing mechanistic insight, indicating that *in vivo* this function is mediated by acetylation of H3K27, a target commonly regulated by CBP/nej and HDAC1. Notably, cells carrying the H3K27R mutation exhibit derepression of Hox genes, closely resembling the phenotype associated with loss-of-function mutations in PRC1 and PRC2 complexes(Pengelly et al., 2013, Classen et al., 2009, Beuchle et al., 2001). However, H3K27R cells also display a pronounced, albeit not complete, reduction in overall H3K27ac levels – in contrast to the nearly complete loss of H3K27ac observed after 24 h of CBP/nej RNAi-expression. This suggests that CBP/nej-mediated acetylation of K27 may occur on both H3 and H3.3 variants, with the anti-H3K27ac antibody likely recognizing both forms and the broad substrate specificity of CBP/nej likely targeting both histone variants. Importantly, acetylation of H3 and H3.3 at K27 may carry distinct functional implications. H3.3 variants appear to be particularly critical for enhancer-associated H3K27ac thereby serving roles in interphase cells (Trovato et al., 2020), whereas acetylation of canonical H3 may regulate chromatin function and DNA replication in S-phase. Indeed, associations of H3K27ac with Origin Recognition Complex-binding sites and its requirement for gene amplification during endoreplication (Aggarwal and Calvi, 2004, Eaton et al., 2011, Hua and Orr-Weaver, 2017, McConnell et al., 2012), suggests that elevated levels of H3K27ac may be essential to establish an open chromatin environment necessary for efficient DNA replication. Collectively, these findings position H3K27ac as a central regulator integrating chromatin structure, DNA replication and genome integrity.

In conclusion, our findings establish a robust mechanistic link between cell cycle dynamics, metabolic signaling and histone modification of H3K27 and H3K9, and specifically implicate H3K27ac as critical mediator of replication fidelity and chromatin organization. These insights highlight the necessity of maintaining balanced histone modification levels to preserve genomic stability, particularly under varying proliferative and metabolic states, with broad implications for developmental biology, aging, and cancer biology.

## EXPERIMENTAL PROCEDURES

### Drosophila genetics

All experiments were performed on *Drosophila melanogaster*. Fly strains (see Table S1) were maintained on standard fly food (10L water, 74,5g agar, 243g dry yeast, 580g corn flour, 552ml molasses, 20.7g Nipagin, 35ml propionic acid) at 18°C – 22°C. Larvae from experimental crosses were allowed to be fed on Bloomington formulation (175.7g Nutry-Fly,1100ml water 20g dry yeast, 1.45g Nipagin in 15ml Ethanol, 4.8ml Propionic acid) and raised at 18°C or 30°C to control GAL80ts-dependent induction of GAL4/UAS. To drive expression of UAS-constructs, such as *UAS-Cdk1-RNAi*, under the control of *rn-GAL4* in the wing pouch, experiments were conducted following protocols from Smith-Bolton et al. (2009), La Fortezza et al. (2016), and Cosolo et al. (2019), with slight modifications. Briefly, larvae with the genotype *rn-GAL4, tub-GAL80^ts^* and appropriate UAS-transgenes were staged by a 6 hours egg collection and raised at 18°C with a density of 50 larvae per vial. Transgene expression was activated by shifting the temperature to 30°C for 24 hours on day 7 after egg deposition (AED). Unless otherwise noted, dissections were performed immediately after this induction period (recovery time point R0). Control genotypes were derived from either rn^ts^ control crosses or sibling animals (+/TM6c). All experiments were performed with at least ≥ 2 independent experimental replicates. Our experimental design did not consider differences between sexes unless for genetic crossing schemes.

### FLP/FRT-Mediated Clone Induction

Mitotic recombination to generate mosaic clones of the *Drosophila* histone gene cluster (ΔHisC) was induced using the FLP/FRT system. Adult flies were allowed to lay eggs for 72 hours at 25 °C. FLP recombinase expression was triggered by a 1-hour heat shock at 37 °C. Larvae were dissected either 48 or 72 hours after heat shock for subsequent analysis. Transgene cassettes providing a total of 12 copies of the Histone gene unit with a H3K27R point mutation (12×His-GUH3-K27R) rescues cell viability of (ΔHisC) mutant clones (see (Pengelly et al., 2013)).

### Immunohistochemistry of wing imaginal discs

To visualize histone modifications, wing discs from third instar larvae were dissected and fixed for 10 minutes at room temperature in 4% paraformaldehyde in PBS. Washes were performed in PBS containing 0.1% TritonX-100 (PBT). The discs were then incubated with primary antibodies (**Table S1**) in 0.1% PBT, gently mixing 2-overnights at 4°C. During incubation with cross-absorbed secondary antibodies coupled to Alexa Fluorophores (Table S1) at 4°C for 1-overnight, tissues were counterstained with DAPI (0.25 ng/µL, Sigma, D9542). Tissues were mounted using SlowFade Gold Antifade (Invitrogen, S36936). To visualize DNA damage using the anti-H2AvD phosphoS137 antibody, wing discs were fixed 5 minutes at room temperature in 4% paraformaldehyde in PBS. Washes were performed in PBS containing 0.5% TritonX-100. The primary antibody was incubated in 0.5% PBT overnight at 4°C and the secondary antibody in 0.5% PBT for 2 hours at room temperature. For all other staining protocols, discs were fixed for 15 minutes at room temperature in 4% paraformaldehyde in PBS. All washes and incubations were performed in PBS containing 0.1% TritonX-100 as described above.To maintain consistency in staining across genotypes, control and experimental wing discs were processed together in the same vial throughout the protocol and mounted on the same slide whenever possible. Images were acquired using the Leica TCS SP8 Microscope, using the same confocal settings for linked samples and processed using tools in Fiji. Figure panels were assembled in Affinity Designer 2.

### EdU labelling

EdU incorporation was performed using the Click-iT Plus EdU Alexa Fluor 647 Imaging Kit (Table S1). Briefly, larval cuticles were inverted in Schneider’s medium and incubated with EdU (10µM final concentration) at RT for 15 minutes. Cuticles were then fixed in 4% PFA/PBS for 15 minutes, washed for 30 minutes in PBT 0.5%. EdU-Click-iT labeling was performed according to manufacturer’s guidelines. Tissues were washed in PBT 0.1%, after which immunohistochemistry, sample processing and imaging were carried out as described above.

### SA-β-Gal staining

CellEvent senescence detection kit from Invitrogen (Table S1) was used to check senescence-associated β-galactosidase activity, following the manufacturer instructions. Briefly, larvae were dissected in PBS, fixed with 4% PFA, washed with 1% BSA (in PBS), and then incubated in working solution for 2h at 37°C. Washing steps were performed in PBS and PBS containing 0.1% TritonX-100 (PBT). Tissues were counterstained with DAPI (0.25 ng/μl). Further immunohistochemistry analysis and sample mounting was performed as described above.

### Protein synthesis assay using OPP-Click-iT staining

OPP Assays were performed using Click-iT® Plus OPP Protein Synthesis Assay Kits (Invitrogen Molecular Probe) according to manufacturer’s instructions. Briefly, larvae were dissected and inverted cuticles were incubated with a 1:1000 dilution of Component A in Schneider’s medium for 15 minutes on a nutator. Larval cuticles were fixed with 4% paraformaldehyde for 15 minutes, rinsed twice in 0.1% PBT, and permeabilized with 0.5% PBT for 15 minutes. The cuticles were then stained with the Click-iT® cocktail for 30 minutes at room temperature, protected from light. Further immunohistochemistry analysis and sample mounting was performed as described above.

### CUT&Tag sample preparation

Briefly, for each genotype around 80-100 wing discs were dissected in ice cold Schneider’s medium, collected, snap frozen and stored at −80°C until further use. Nuclei were isolated with the help of a mechanical pestle and processed for CUT&Tag analysis as previously described (Zenk et al., 2023, Cardamone et al., 2025). For normalizing the levels of histone H3 modifications and as a proxy of the starting material in each condition, we also generated samples for total histone H3. Libraries were prepared by supplementing the NEBNext HighFidelity 2× PCR Master Mix with 1 pg of Tn5-tagmented lambda DNA (New England Biolabs, #N3011S) as a spike-in normalizer. The library indexing was performed using Illumina i5 and i7 (Buenrostro et al., 2015) through 15 cycles (1 x 5 min at 72 °C, 1 x 30 sec at 98°C, 13 x 10 s at 98 °C, 30 sec at 63 °C, 1 x 1 min at 72 °C, hold at 4 °C). The libraries were purified using Nucleomag NGS Clean-up and Size Select beads (Macherey-Nagel, #744970.50). Quality checks were done with Qbit DNA HS Assay (ThermoScientific, #Q32854) and Bioanalyzer (Agilent). Finally, two independent library replicates were sent for next generation sequencing.

## QUANTIFICATIONS, BIOINFORMATIC ANALYSIS AND STATISTICS

### General comments

For all quantifications of imaging data, control and experimental samples were processed together and imaged in parallel, using the same confocal settings. Images were processed, analyzed and quantified using tools in Fiji (ImageJ 2.9.0) (Schindelin et al., 2012). Extreme care was taken to apply consistent methods (i.e., number of projected sections, thresholding methods, processing) for image analysis. Figure panels were assembled using Affinity Designer 2.6.2. Statistical analyses were performed in Graphpad Prism. Illustrations were created in Biorender.

### Quantification of histone modifications

For quantification of levels of histone modifications, an xy-representation containing the maximum number of pouch and hinge cells was generated using a sum projection of the relevant channel. Three square regions of interest (ROIs) measuring 15 × 15 μm were placed in distinct areas of either the pouch or the hinge, carefully excluding regions with developmentally arrested cells. The mean intensity of the modification signal was measured within each ROI. For each disc, the average pouch signal was normalized to the average hinge signal and reported as a pouch-to-hinge intensity ratio.

### Quantification of EdU incorporation

Figure S1.1: To first create a nuclear mask, an xy-representation containing the maximum number of pouch nuclei was generated using a sum projection of the DAPI channel. The background signal was reduced using the “Subtract Background” function with a rolling ball radius of 100 pixels. Local contrast was enhanced with the “Enhance Local Contrast (CLAHE)” function, using a block size of 127, 256 histogram bins, and a maximum slope of 3. Automatic thresholding was performed using the “Moments” method with the “dark” option selected, and the image was then converted into a binary mask of DAPI-positive nuclear areas. A nuclear ROI was generated from this mask using the selection tool. The pouch region was manually selected based on wing fold morphology. To isolate the nuclei within the pouch, a binary “AND” operation was used to combine nuclear mask and pouch selection. This final pouch-specific DAPI mask was used to measure EdU signal intensity.

Figure 6: To create an EdU mask, an xy-representation containing the maximum number of pouch nuclei was generated using a sum projection of the EdU channel. A “Gaussian blur” (σ = 2) was applied to reduce noise and smooth the signal. Automatic thresholding was then performed using the “Moments” method with the “dark” option selected. The image was converted into a binary EdU mask with a black background. A ROI was generated from this mask using the selection tool to define EdU-positive pixels. Separately, the pouch region was manually selected based on wing fold morphology. To isolate EdU-positive pixels specifically within the pouch, a binary “AND” operation was applied to combine the EdU selection with the pouch selection. This final pouch-specific EdU mask was used to measure EdU signal intensity.

To assess changes in the number of proliferating cells, pouch-specific DAPI and EdU masks were generated as described above. A binary “AND” operation was used to extract EdU-positive nuclei located within DAPI-positive regions in the pouch. The area of EdU-positive nuclei was then divided by the total area of EdU and DAPI-positive regions within the pouch to calculate the fraction of proliferating cells. This EdU-to-DAPI area ratio was calculated per disc and used for comparative analysis.

### Quantification of OPP levels in wing imaginal discs

An xy-representation containing the maximum number of pouch cells was generated using a sum projection of the relevant channels. In both control and constitutively active insulin receptor-expressing discs, the pouch region was manually selected based on wing fold morphology. Mean OPP intensity was then measured within the defined pouch region for each disc.

### Quantification of pouch size

For both control and constitutively active insulin receptor-expressing discs, a slice containing the maximum number of pouch cells was selected. In each case, the pouch region was manually defined based on wing fold morphology. The area size of the selected pouch region was then measured for each disc.

### Cell cycle phase classification using FUCCI and quantifications of H3, Total Acetyl Lysine, H3K27ac, H3K18ac and H4K8ac level

To classify nuclei in the wing pouch according to cell cycle phase and to quantitatively correlate levels of H3 and acetylation levels with each phase, a custom image analysis workflow was implemented in ImageJ/Fiji. A single z-slice containing the highest number of pouch cells was selected from each control disc. The image stack was split into individual channels (e.g., DAPI, GFP-E2f1^1-230^, mRFP-NLS-CycB^1-266^, H3K27ac). To generate a nuclear mask, the DAPI, GFP, RFP, and other nuclear markers were averaged sequentially using the Image Calculator. The resulting image was thresholded using the Otsu method (upper limit set to 255) and converted into a binary mask defining nuclear pixels. A nuclear pixel selection was generated from the binary image and added to the ROI Manager. The pouch region was manually defined based on wing fold morphology. A binary “AND” operation was used to restrict the nuclear pixel mask to the pouch, producing a final pouch-specific nuclear mask. This mask was overlaid on the GFP, RFP, and other nuclear marker channels, and the X/Y coordinates of each ROI were used to extract signal intensities for each nuclear pixel. Threshold values for GFP and RFP determined based on average intensity of 5 nuclei in early S-phase. Each nuclear pixel was classified into a cell cycle phase using these thresholds, following FUCCI-based criteria: G2 phase: GFP > threshold, RFP > threshold; G1 phase: GFP > threshold, RFP ≤ threshold; Early/mid S phase: GFP ≤ threshold, RFP ≤ threshold; Late S phase: GFP ≤ threshold, RFP > threshold. For each nuclear pixel, the intensity of the corresponding nuclear markers (e.g., H3, acetyl-lysine, H3K27ac) was recorded. To account for inter-sample variability, intensity values within each cell cycle phase were normalized by dividing them by the total nuclear marker signal summed across all phases (G1, early S, late S, and G2) within the same disc. This normalization provided the relative contribution of each cell cycle phase to the total signal, allowing for direct comparisons across samples.

### γH2Av intensity quantifications

To create a nuclear mask, an xy-representation containing the maximum number of pouch nuclei was generated using a sum projection of the DAPI channel. The background signal was reduced using the “Subtract Background” function with a rolling ball radius of 100 pixels. Local contrast was enhanced with the “Enhance Local Contrast (CLAHE)” function, using a block size of 127, 256 histogram bins, and a maximum slope of 3. Automatic thresholding was performed using the “Moments” method with the “dark” option selected, and the image was then converted into a binary mask. A nuclear ROI was generated from this mask using the selection tool. The pouch region was manually selected based on wing fold morphology. To isolate the nuclear pixels within the pouch, a binary “AND” operation was used to combine nuclear mask and pouch selection. This final mask was used to measure γH2Av and DAPI intensities for each disc. γH2Av intensities were normalized to the corresponding DAPI intensity.

### CUT&Tag Bioinformatic Analysis

The CUT&Tag data processing was performed as described before with some modifications (Atinbayeva et al., 2024, Ibarra-Morales et al., 2021, Cardamone et al., 2025). As our quantitative CUT&Tag sequencing libraries contained DNA from *D. melanogaster* and spike-in of *lambda* phage genome (Genbank: J02459.1), we mapped the raw paired-end reads to a constructed hybrid dm6 and *Lambda* phage genome using *snakePipes-v3.0.0 DNA-mapping* (Bhardwaj et al., 2019) with updated Bowtie2 mapping parameters, as well as adapter trimming and MAPQ ≥ 1 filtering (*DNAmapping --mapq 1 --trim --fastqc --properPairs -- dedup --cutntag --DAG --alignerOpts=’--local --very-sensitive-local --no-discordant --no-mixed -I 10 -X 700’*). To assess reproducibility of both replicates, we utilised the *multiBamSummary* and *plotCorrelation* from *deepTools-v.2.5.7* to compute the Spearman correlation between the two replicates. Aligned replicates were merged before normalization to both H3 and spike-in signals. The code used for the H3 and *Lambda* normalization analysis was based on work done by Yinxiu Zhan (https://github.com/zhanyinx/atinbayeva_paper_2023) for (Atinbayeva et al., 2024). CUT&Tag peaks were called using *snakePipes-v3.0.0 ChIPseq* with *MACS2-v 2.2.9.1* (*ChIPseq -d Output_spikein hybrid.yaml H3_spikein_config.yaml --peakCaller MACS2 --cutntag --peakCallerOptions “--broad --qvalue 0.5 –f BAMPE” --DAG*). Normalized bigWig and bam files were generated with *deepTools-v3.5.6* and used to compute matrices of signal enrichment +/− 5kb around transcription start sites (TSS) or peak centers of shared or unique peaks for *wt* and *Cdk1KD*. Coverage heatmaps and profiles were created using *plotHeatmap* or *plotProfile* from *deepTools-v.2.5.7*.

For the quantification, the number of PE reads were counted in each shared or unique peak for *wt* and *Cdk1KD* (*multiBamSummary BED-file --bamfiles* <*input files*> *--extendReads --outRawCounts* <*file*>) as well as in 500 bp bins across the dm6 genome (*multiBamSummary bins --bamfiles* <*input files*> *-binSize 500 --extendReads --outRawCounts* <*file*>). The *DESeq2-v1.38.3* analysis was performed using the previously computed H3 and spike-in scaling factors from merged data for size factor. Bins and peaks with basemean (average read count across samples) of less than or equal to 25 read counts were discarded. The design matrix was setup to compare the samples by condition and correct for replicate effects (*design* = *∼replicate* + *condition*). Finally, we executed the *DESeq2* shrinkage of log2 fold changes (*type* = *‘normal’*).

## CODE AND DATA AVAILABILITY

All data, workflows and FIJI based algorithms necessary to interpret the imaging data are included within the manuscript. Original imaging data and any additional information required to reanalyze the data reported in this paper will be shared by AKC upon request. The code used for the mapping, normalization, peak calling and signal quantification is available on Github: https://github.com/LaraH9/nogay_et_al_2025. The CUT&Tag sequencing data generated in this study is provided on NCBI SRA: https://www.ncbi.nlm.nih.gov/sra/PRJNA1300380.

## AUTHOR CONTRIBUTIONS

Conceptualization LN, AKC; Data curation LN, LH, AKC; Formal analysis LN, LH, AKC; Funding acquisition AKC; Investigation LN, AVM; Methodology LN, AVM, LH, FC, IG, AS, AF, LE, AKC; Project administration IG, AKC; Supervision NI, AKC; Validation LN, AVM, FC, IG; Visualization LN, LH, AKC; Writing – original draft LN, AKC; Writing – review & editing LN, AVM, LH, AKC

## ACKNOWLEDGEMENTS

We thank the staff of the Life Imaging Center (LIC) in the Hilde Mangold House (HMH) of the Albert-Ludwigs-University of Freiburg for help with their confocal microscopy resources, and the excellent support in image recording. We specifically thank the DFG for supporting our imaging work through project number 414136422. We thank the Bioinformatics and Sequencing facilities at the MPI-IE for supporting this work. We thank Galina Erikson for supporting our bioinformatic analysis. We thank Laura Bauttita, Melissa Harrison, Iswar Hariharan, Dirk Bohmann and Jürg Müller for sharing reagents and the Bloomington *Drosophila* Stock Center (BDSC), the Vienna *Drosophila* Stock Collection (VDRC), the University of Zurich ORFeome Project (FlyORF) and the Developmental Studies Hybridoma Bank (DSHB) for providing fly stocks and antibodies. We thank the IMPRS-EBM and SGBM graduate schools for supporting our doctoral researchers.

## SUPPLEMENTARY FIGURES

**Figure S1.1.**
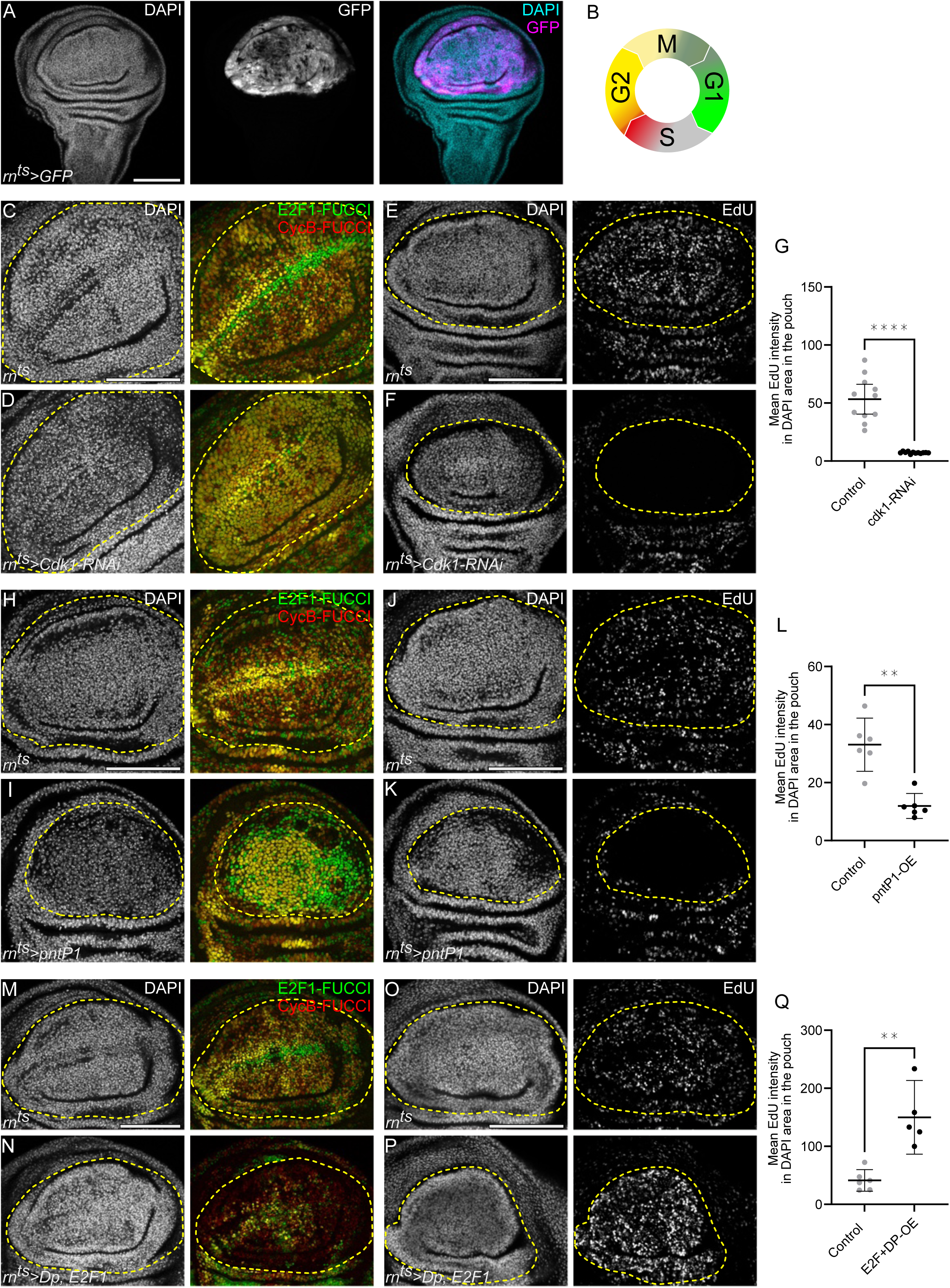
Cell cycle manipulation models for assessing histone modification dynamics **A.** A control wing disc after 24 h of *UAS-GFP*-expression in the pouch, under the control of the *rn-GAL4* (*rotund-GAL4*) driver. GFP-expression visualizes the tissue domain subject to manipulation throughout this study (magenta). **B.** Schematic representation of the Fly-FUCCI cell cycle reporter system, utilizing the degradable *GFP-E2F1*^1-230^ (green) and *mRFP-NLS-CycB*^1-266^ (red) as sensors to visualize cell cycle phases. Individual fluorophore expression indicates cells in G1 or late S-phase, respectively. Combined expression of both GFP and RFP labels cells is observed in early and late G2. Cells in late mitosis and early S-phase lack expression of either GFP or RFP. Cells in early S-phase are specifically detected by EdU incorporation (Crucianelli et al., 2022; Zielke et al., 2014). Cells in mitosis are positioned in very apical positions of the tissue and are not represented in our study. **C-G.** Cell cycle dynamics in control discs **(C, E)** and discs expressing *Cdk1-RNAi* for 24 h under the control of *rn-GAL4* **(D, F)**. FUCCI reporters *GFP-E2F1*^1-230^ (green) and *mRFP-NLS-CycB*^1-266^ (red) were used to visualize cell cycle phases **(C, D)** and EdU incorporation was used to detect DNA replication in S-phase cells **(E, F).** Cdk1 knockdown in the wing pouch domain leads to a cell cycle phase shift toward G2 phase **(D)**, and a corresponding loss of EdU incorporation **(F)** combined demonstrating a pronounced arrest in G2. **(G)** Quantification of mean EdU intensity per DAPI area in the pouch region of control and *Cdk1-RNAi*-expressing discs, serving as a proxy for relative DNA replication activity. Mean and 95% CI are shown. Statistical significance was tested using two-tailed Welch’s test, p value <0.0001 (control discs: n=11, *Cdk1-RNAi*-expressing discs: n=12). **H-L.** Cell cycle dynamics in control discs **(H, J)** and discs expressing *pntP1* **(I, K)** for 24 h under the control of *rn-GAL4.* FUCCI reporters *GFP-E2F1*^1-230^ (green) and *mRFP-NLS-CycB*^1-266^ (red) were used to visualize cell cycle phases **(H, I)** and EdU incorporation was used to detect DNA replication in S-phase cells **(J, K).** Pointed-P1 overexpression in the wing pouch domain leads to a cell cycle phase shift towards either the G1 or the G2 phase **(I)**, correlating with a loss of EdU incorporation **(K)**. Combined, this demonstrates a pronounced arrest in either G1 or G2. Mechanistically, the choice for either cell cycle phase is not known. **(L)** Quantification of mean EdU intensity per DAPI area in the pouch region of control and *pntP1*-expressing discs, serving as a proxy for relative DNA replication activity. Mean and 95% CI is shown. Statistical significance was tested using two-tailed Mann-Whitney test, p value=0.0043 (control discs: n=6, *pntP1*-expressing discs: n=6). **M-Q.** Cell cycle dynamics in control discs **(M, O)** and discs expressing *Dp, E2F1* **(N, P)** for 24 h under the control of *rn-GAL4.* FUCCI reporters *GFP-E2F1*^1-230^ (green) and *mRFP-NLS-CycB*^1-266^ (red) were used to visualize cell cycle phases **(M, N)** and EdU incorporation was used to detect DNA replication in S-phase cells **(O,P).** dDp and dE2F1 co-expression in the wing pouch domain reduces the number of cells in either G1 or G2 **(N)**, and an increase in S-phase cells as well as an elevation of EdU incorporation **(P)** in the wing pouch region, confirming frequent entry and rapid progression through the cell cycle. **(Q)** Quantification of mean EdU intensity per DAPI area in the pouch region of control and *Dp, E2F1*-coexpressing discs, serving as a proxy for relative DNA replication activity. Mean and 95% CI is shown. Statistical significance was tested using two-tailed Welch’s t test, p value=0.0069 (control discs: n=6, *Dp, E2F1*-expressing discs: n=5). Discs were stained with DAPI to visualize nuclei. Fluorescence intensities are reported as arbitrary units. Yellow dashed lines represent the pouch region of wing discs. Sum projections of multiple confocal sections are shown in **A, E-F, J-K, M-N** and **O-P.** Maximum projections of multiple confocal sections are shown in **C-D** and **H-I.** Scale bars: 100 μm.

**Figure S1.2.**
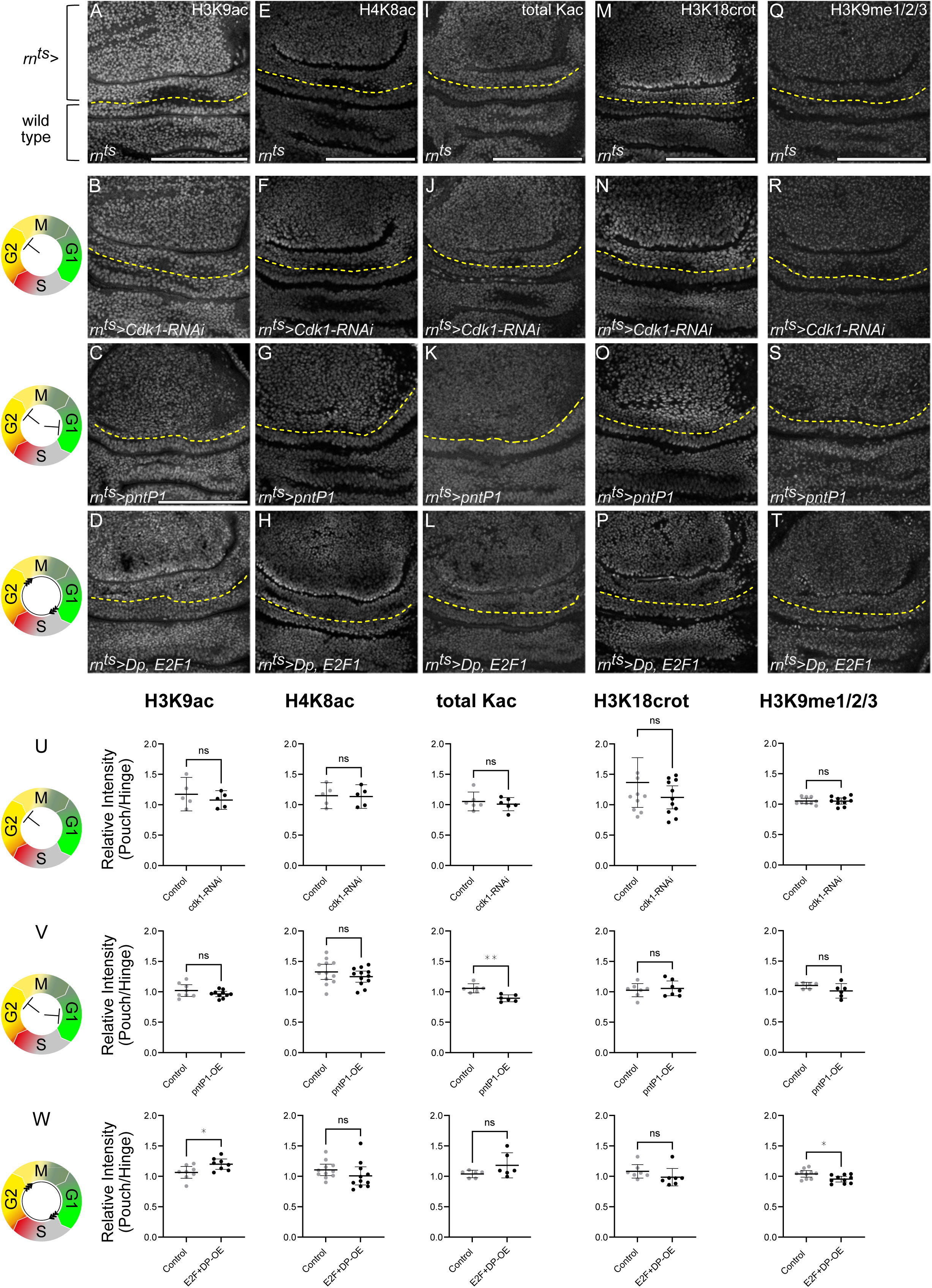
Histone modifications are largely insensitive to cell cycle manipulations **A-D.** Immunostaining for H3K9ac in control **(A)**, *Cdk1-RNAi*-expressing **(B)**, *pntP1*-expressing **(C)** and *Dp, E2F1*-expressing **(D)** discs. Expression was induced for 24 h in the wing pouch using *rn-GAL4*. The *rn-GAL4* expression pouch domain is represented by the tissue above the yellow line (rn^ts^>). The tissue below the yellow line represents wild type tissues of the hinge. Icons represent the experimentally verified FUCCI cell cycle status (see **Fig S1.1**) in each condition: G2-phase arrest in *Cdk1-RNAi*-expressing discs **(B)**, combined G1 and G2-phase arrest in *pntP1*-expressing discs **(C)**, and cell cycle acceleration in *Dp, E2F1*-coexpressing discs **(D)**. **E-H**. Immunostaining for H4K8ac in control **(E)**, *Cdk1-RNAi*-expressing **(F)**, *pntP1*-expressing **(G)** and *Dp, E2F1*-coexpressing **(H)** discs. **I-L.** Immunostaining for total acetylated lysines in control **(I)**, *Cdk1-RNAi*-expressing **(J)**, *pntP1*-expressing **(K)** and *Dp, E2F1*-coexpressing **(L)** discs. **M-P.** Immunostaining for H3K18crotonylation in control **(M)**, *Cdk1-RNAi*-expressing **(N)**, *pntP1*-expressing **(O)** and *Dp, E2F1*-coexpressing **(P)** discs. **Q-T.** Immunostaining for total H3K9me1/2/3 methylation in control **(I)**, *Cdk1-RNAi*-expressing **(J)**, *pntP1*-expressing **(K)** and *Dp, E2F1*-coexpressing **(L)** discs. **U.** Quantification of relative signal intensities for H3K9ac, H4K8ac, total acetylated lysine, H3K18crotonylation, and H3K9me1/2/3, presented as pouch-to-hinge ratios (rn^ts^-to-wild-type-cell ratio) in *Cdk1-RNAi*-expressing discs. Mean and 95% CI is shown. Statistical significance was tested using two-tailed Unpaired t test, p value= 0.4268 for H3K9ac (control discs: n=5, *Cdk1-RNAi*-expressing discs: n=5); two-tailed Unpaired t test, p value=0.8936 for H4K8ac (control discs: n=5, *Cdk1-RNAi*-expressing discs: n=5); two-tailed Unpaired t test, p value=0.5553 for total acetylated lysines (control discs: n=6, *Cdk1-RNAi*-expressing discs: n=6); two-tailed Mann-Whitney test, p value=0.4779 for H3K18crot (control discs: n=11, *Cdk1-RNAi*-expressing discs: n=11); two-tailed Unpaired t test, p value=0.9619 for H3K9me1/2/3 (control discs: n=9, *Cdk1-RNAi*-expressing discs: n=10). **V.** Quantification of relative signal intensities for H3K9ac, H4K8ac, total acetylated lysine, H3K18crotonylation, and H3K9me1/2/3, presented as pouch-to-hinge ratios (rn^ts^-to-wild-type-cell ratio) in *pntP1*-expressing discs. Mean and 95% CI is shown. Statistical significance was tested using two-tailed Unpaired t test, p value= 0.1975 for H3K9ac (control discs: n=8, *pntP1*-expressing discs: n=10); two-tailed Unpaired t test, p value=0.2752 for H4K8ac (control discs: n=12, *pntP1*-expressing discs: n=12); two-tailed Unpaired t test, p value=0.0012 for total acetylated lysines (control discs: n=6, *pntP1*-expressing discs: n=6); two-tailed Unpaired t test, p value=0.6621 for H3K18crot (control discs: n=7, *pntP1*-expressing discs: n=7); two-tailed Mann-Whitney test, p value=0.2403 for H3K9me1/2/3 (control discs: n=6, *pntP1*-expressing discs: n=6). **W.** Quantification of relative signal intensities for H3K9ac, H4K8ac, total acetylated lysine, H3K18crotonylation, and H3K9me1/2/3, presented as pouch-to-hinge ratios (rn^ts^-to-wild-type-cell ratio) in *Dp, E2F1*-coexpressing discs. Mean and 95% CI is shown. Statistical significance was tested using two-tailed Unpaired t test, p value= 0.0282 for H3K9ac (control discs: n=8, *Dp, E2F1*-coexpressing discs: n=8); two-tailed Mann-Whitney test, p value=0.0759 for H4K8ac (control discs: n=11, *Dp, E2F1*-coexpressing discs: n=11); two-tailed Mann-Whitney test, p value=0.2403 for total acetylated lysines (control discs: n=6, *Dp, E2F1*-coexpressing discs: n=6); two-tailed Mann-Whitney test, p value=0.0728 for H3K18crot (control discs: n=7, *Dp, E2F1*-coexpressing discs: n=7); two-tailed Unpaired t test, p value=0.0154 for H3K9me1/2/3 (control discs: n=10, *Dp, E2F1*-coexpressing discs: n=10). Fluorescence intensities are reported as arbitrary units. Sum projections of multiple confocal sections are shown in **C-T.** Scale bars: 100 μm. Please refer to **(Fig S1.3B)** for corresponding DAPI images of all discs shown.

**Figure S1.3.**
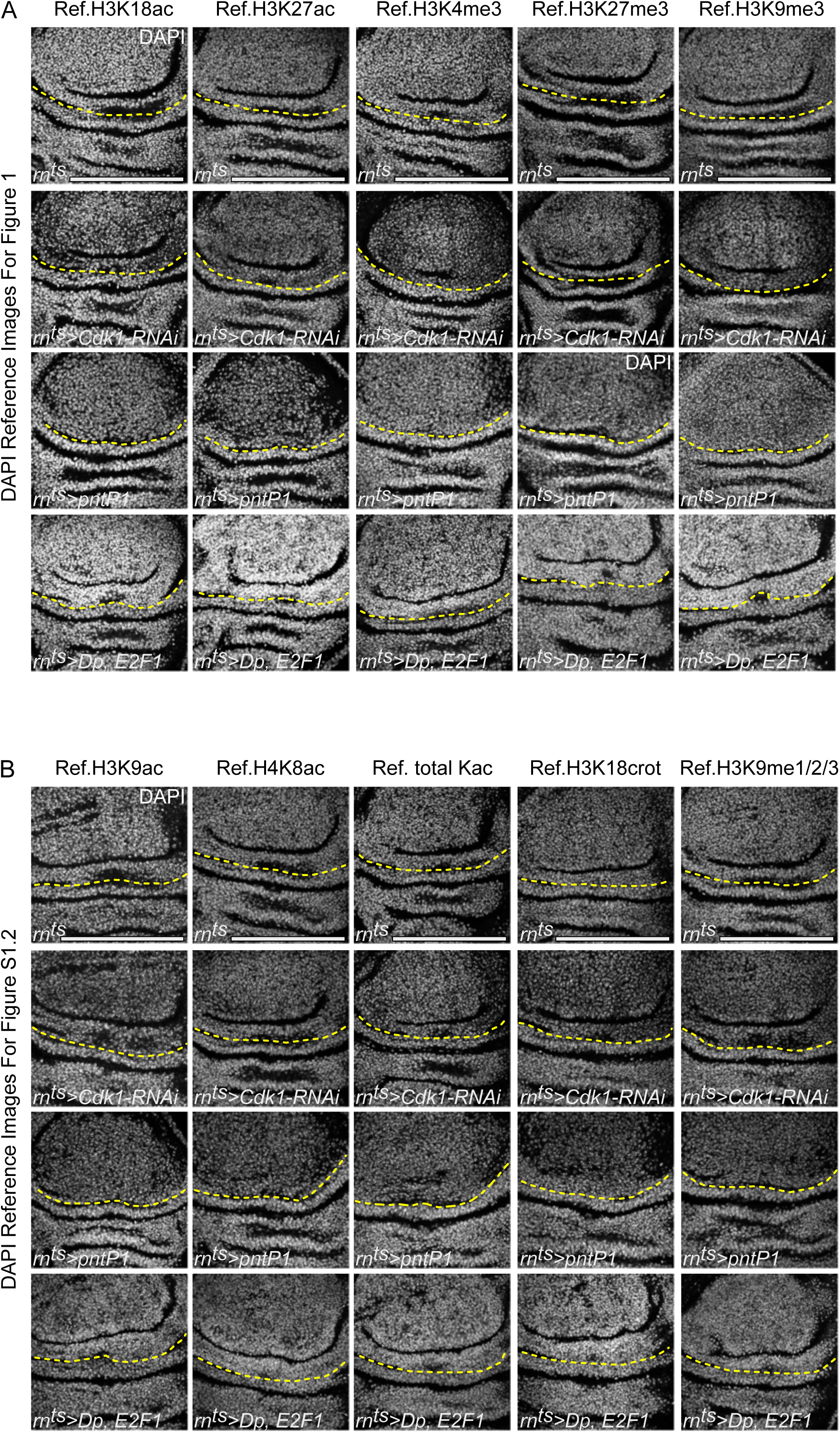
Corresponding DAPI images for Fig 1 and Fig S1.2 Representation of DAPI images matched to the images of histone modifications in control, *Cdk1-RNAi-*expressing, *pntP1-*expressing and *Dp, E2F1*-coexpressing wing discs shown in Fig 1 and Fig S1.2. Discs were stained with DAPI to visualize nuclei. Sum and max projections of multiple confocal sections are shown. Scale bars: 100 μm.

**Figure S2.1.**
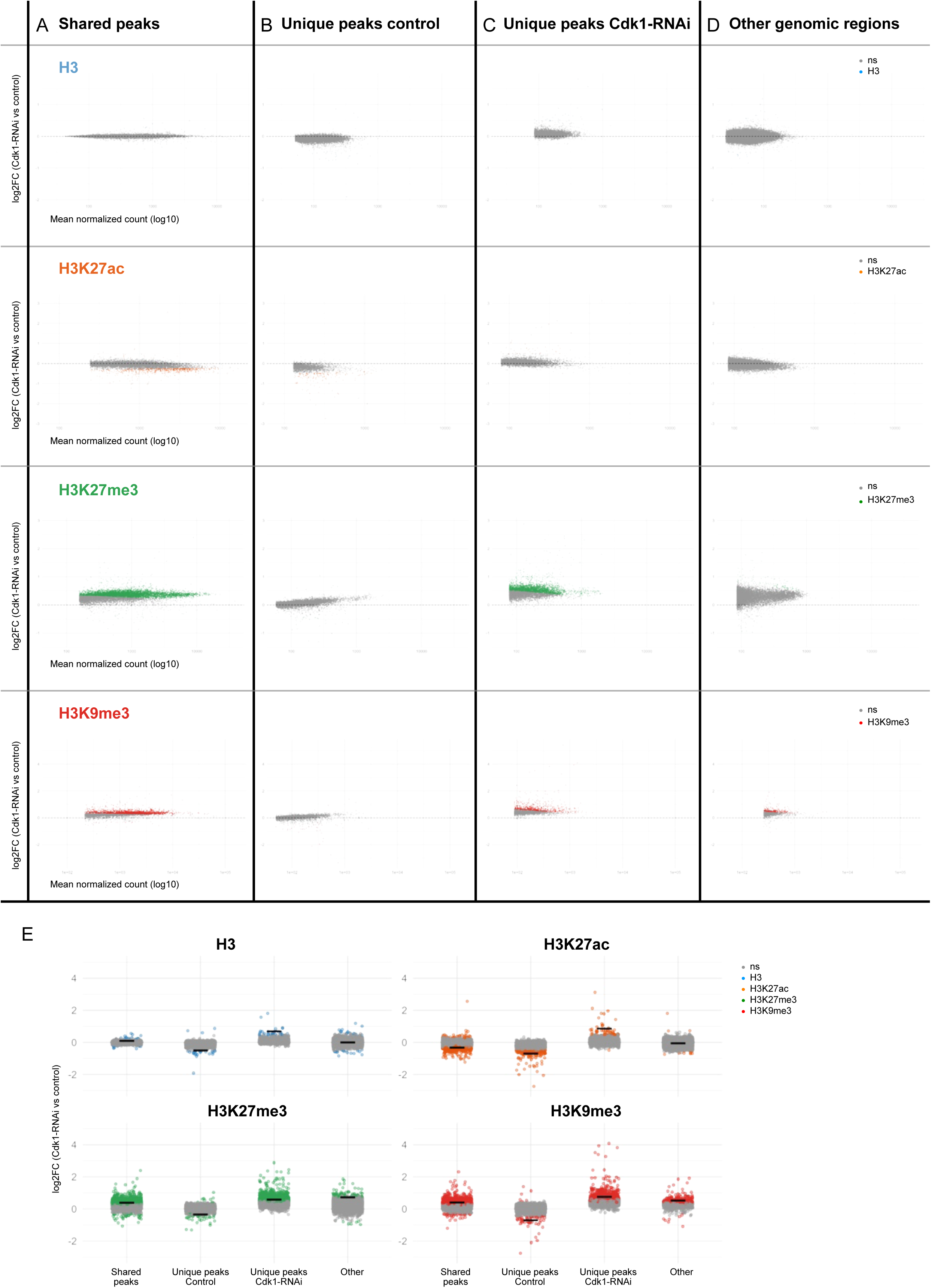
Comparative analysis of Histone H3 and H3K27ac, H3K27me3, and H3K9me3 histone modifications by CUT&Tag in control and Cdk1 RNAi-expressing discs **A-D.** MA plots displaying differential enrichment of histone modifications analysed by by CUT&Tag for H3, H3K27ac, H3K27me3, and H3K9me3 (in rows). Each point represents a shared or unique genomic peak identified by our analysis in control and Cdk1-RNAi expressing discs (shared peaks **(A)**, unique peaks in control **(B)**, unique peaks in cdk1-RNAi **(C)**), or a 500 bp genomic fragment not called as any peak in our analysis (other genomic regions **(D)**). The x-axis displays the mean normalized read count on a log10 scale, representing an average signal intensity across samples or conditions. The y-axis displays the log2 fold change (log2 ratio of read counts) between the Cdk1-RNAi and control conditions, reflecting the magnitude and direction of differential enrichment. Peaks are color-coded by modification type: blue for H3, orange for H3K27ac, green for H3K27me3, and red for H3K9me3. Regions with an adjusted p-value<0.1 are colored accordingly, highlighting statistically significant differential peaks; non-significant peaks (ns, adjusted p-value ≥0.1) are shown in gray. **E.** Each subplot depicts the distribution of log2 fold change values (Cdk1-RNAi compared to control) for genomic regions categorized as Shared (peaks detected in both conditions), Control-Unique (peaks specific to control), Cdk1-RNAi-Unique (peaks specific to Cdk1-RNAi expressing discs), and Other (genomic fragments of 500bp not included in any called peaks). The x-axis indicates these categorical groups, and the y-axis shows the log2 fold change of normalized counts between Cdk1-RNAi and control samples. Each point represents an individual genomic region, colored by histone mark: blue (H3), orange (H3K27ac), green (H3K27me3), and red (H3K9me3). Black horizontal bars indicate the mean log2 fold change within each group. Peaks are color-coded by modification type: blue for H3, orange for H3K27ac, green for H3K27me3, and red for H3K9me3. Regions with an adjusted p-value<0.1 are colored accordingly, highlighting statistically significant differential peaks; non-significant peaks (ns, adjusted p-value ≥0.1) are shown in gray.

**Figure S2.2.**
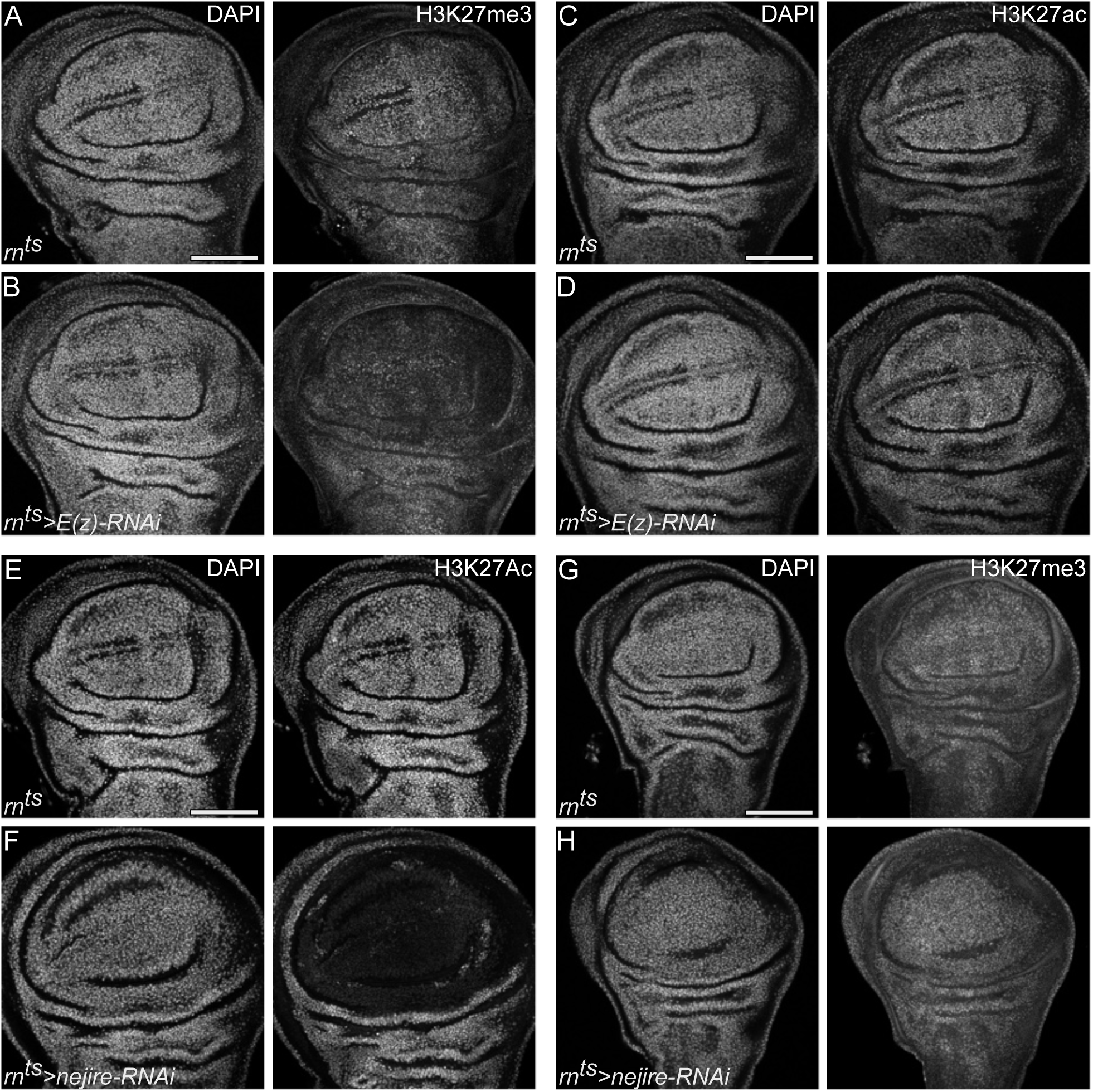
Cell cycle–dependent fluctuations of H3K27ac and H3K27me3 are not due to competitive residue occupancy **A-B.** H3K27me3 staining in control **(A)** or *E(z)-RNAi*-expressing **(B)** discs. Knockdown of the H3K27me3 writer E(z) causes a reduction in H3K27me3 level in the pouch of the wing disc. **C-D.** H3K27ac staining in control **(C)** or *E(z)-RNAi*-expressing **(D)** discs. Knockdown of the H3K27me writer E(z) causes no corresponding increase in H3K27ac level. **E-F.** H3K27ac staining in control **(E)** and *nejire-RNAi*-expressing **(F)** discs. Knockdown of the H3K27ac writer CBP/nej causes a reduction of H3K27ac level in the pouch of the wing disc. **G-H.** H3K27me3 staining in control **(G)** and *nejire-RNAi*-expressing **(H)** discs. Knockdown of the H3K27ac writer CBP/nej causes no corresponding increase in H3K27me3 level. Discs were stained with DAPI to visualize nuclei. Maximum projections of multiple confocal sections are shown in **A-B.** Sum projections of multiple confocal sections are shown in **C-D, E-F** and **G-H.** Scale bars: 100 μm.

**Figure S3.1.**
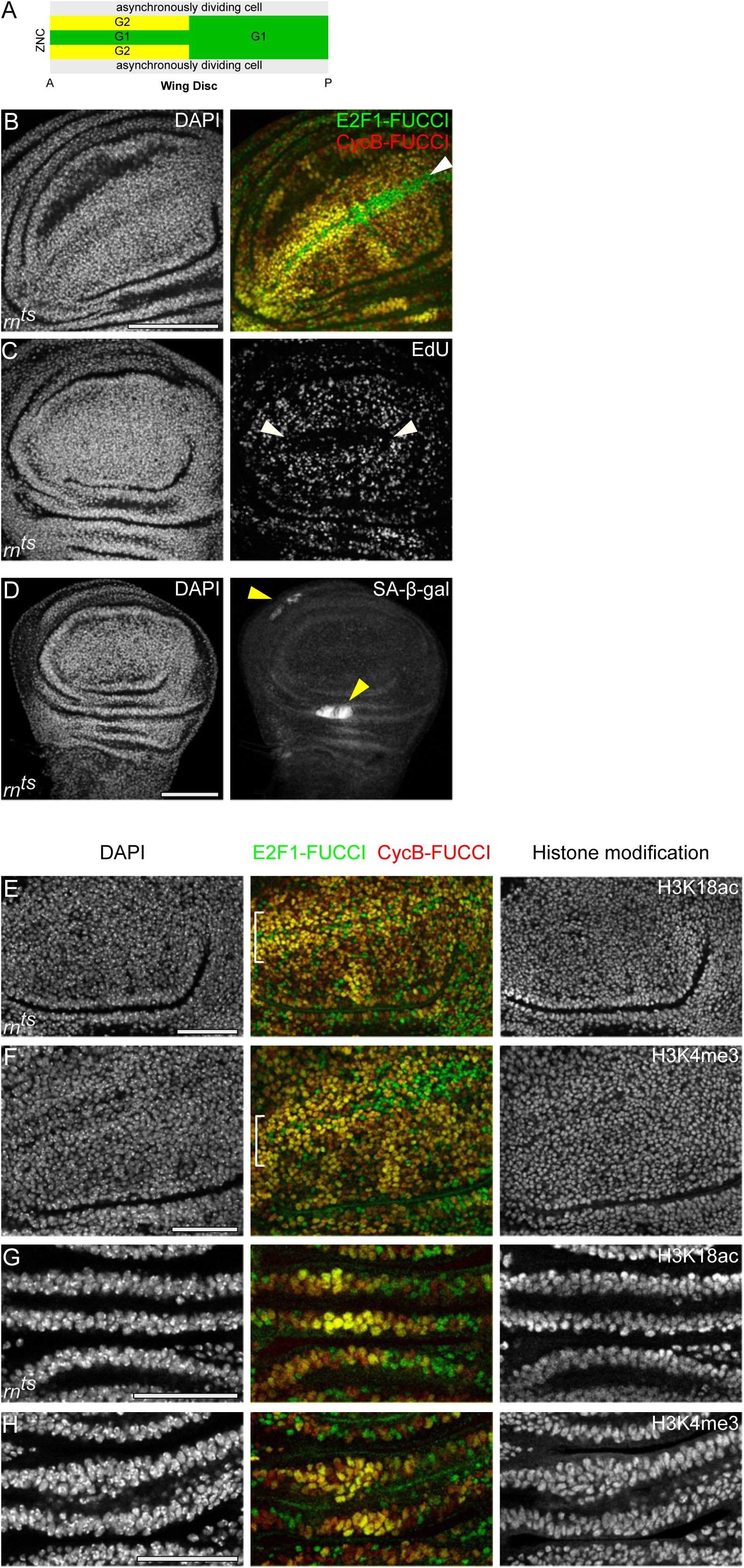
Visualization of cell cycle-arrested cell populations and histone modifications in developing wing imaginal disc **A.** Schematic representation of the zone of non-proliferation cells (ZNC) (adapted from (Johnston and Edgar, 1998)). **B-C.** The Fly-FUCCI system and EdU incorporation assay were used to visualize the spatial pattern of cell cycle arrest within the ZNC. In the anterior ZNC, central margin cells arrest in G1 phase, while adjacent cells arrest in G2. Cells in the posterior ZNC uniformly arrest in G1. The entire ZNC region also lacks EdU incorporation, indicating absence of S-phase entry. White arrowheads indicate the ZNC. **D.** Elevated SA-β-gal activity marks two clusters of cell cycle–arrested cells undergoing programmed senescence (indicated by yellow arrowheads) in developing wing discs. **E,F.** Immunostaining for H3K18ac **(E)** and H3K4me3 **(F)** in the zone of non-proliferation cells (ZNC) of developing wing discs. Neither histone modification shows detectable changes in this region. **G,H.** Immunostaining for H3K18ac **(G)** and H3K4me3 **(H)** in senescent cell populations of the hinge in developing wing discs. Neither histone modification shows detectable changes in this region. Discs were stained with DAPI to visualize nuclei. Maximum projection of multiple confocal sections is shown in **C.** Sum projections of multiple confocal sections are shown in **A-B and D.** Scale bars: 100 μm in **B-D**; 50 μm in **E-H.**

**Figure S3.2.**
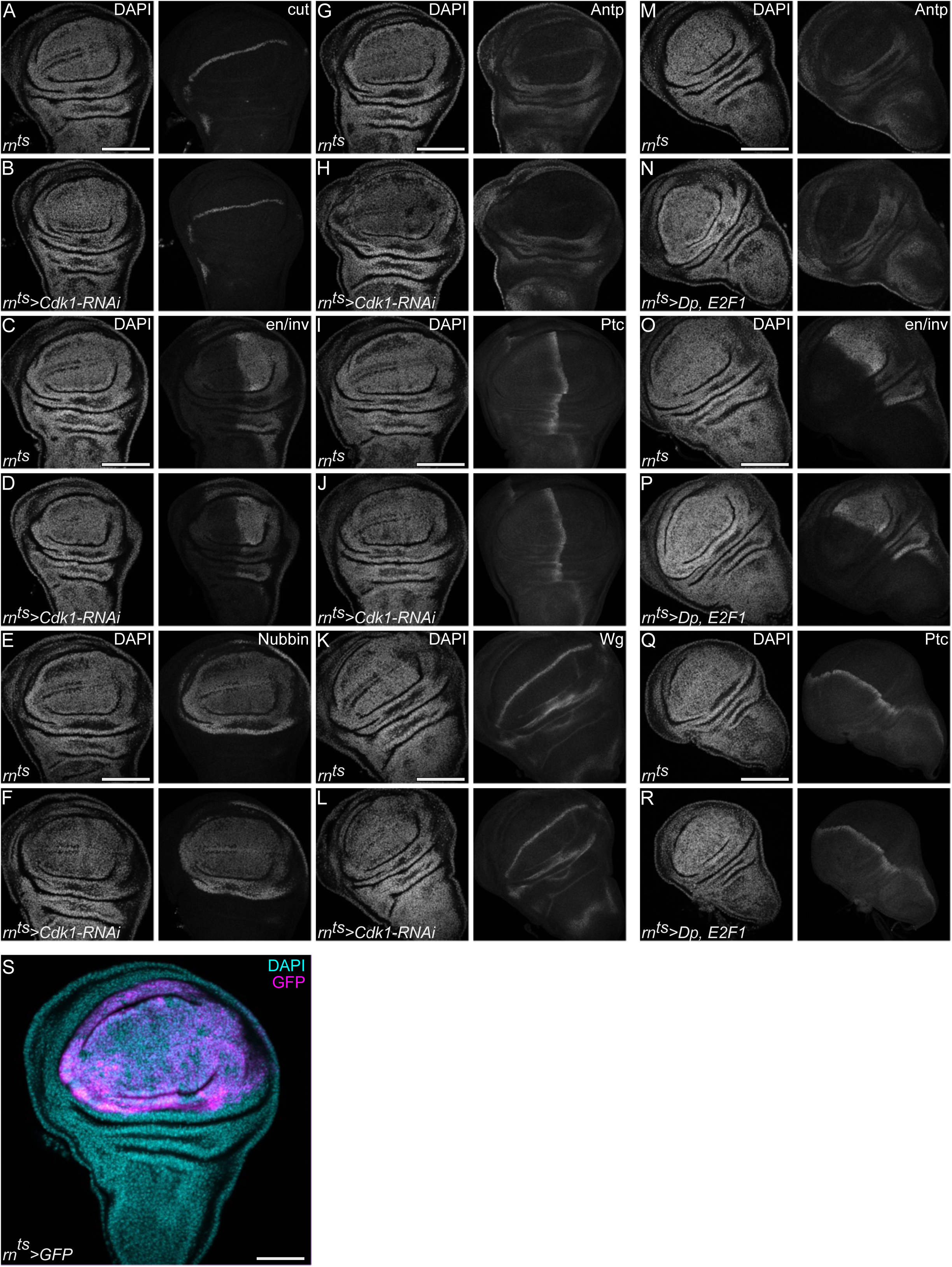
Polycomb target gene expression remains stable despite altered cell cycle dynamics **A-L.** Immunostaining for proteins expressed by Polycomb target genes: Cut **(A, B),** Engrailed/invected **(C, D),** Nubbin **(E, F),** Antennapedia **(G, H),** Patched **(I, J)** and Wingless **(K, L)** in control **(A, C, E, G, I, K)** and *Cdk1-RNAi*-expressing **(B, D, F, H, J, L)** wing discs. **M-R.** Immunostaining for proteins expressed by Polycomb target genes: Antennapedia **(M, N),** Engrailed/i**nvected (O, P)** and Patched **(Q, R)** in control **(M, O, Q)** and *Dp, E2F1*-coexpressing **(N, P, R)** wing discs. **S.** A control wing disc after 24 h of *UAS-GFP*-expression in the pouch (magenta), under the control of the *rn-GAL4* (*rotund-GAL4*) driver, displayed to reference the manipulated pouch domain in Figures A-R. Please note that this is the same disc shown in **Fig S1.1 A**. Discs were stained with DAPI to visualize nuclei. Sum projections of multiple confocal sections are shown in **A-S.** Scale bars: 100 μm.

**Figure S4.1.**
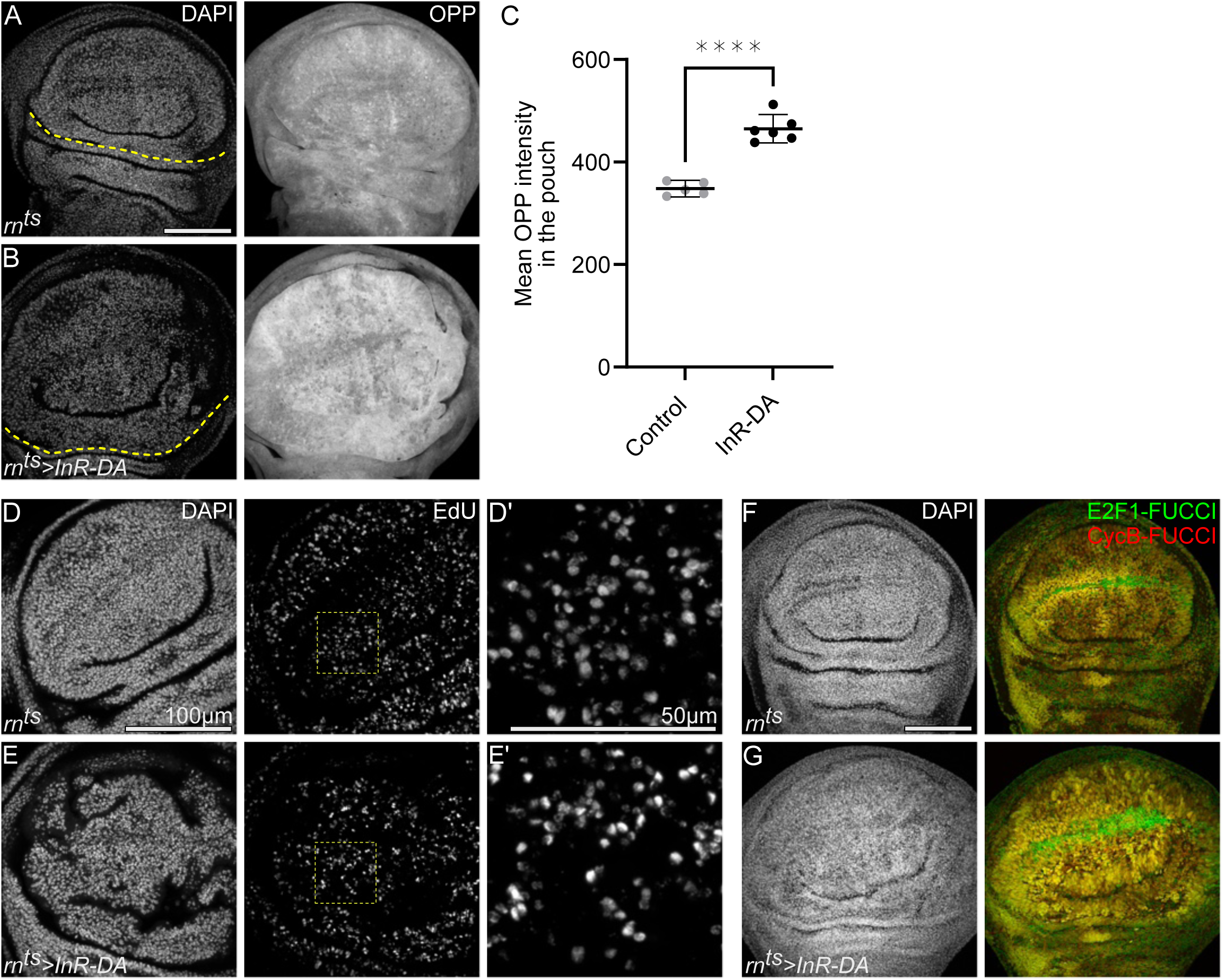
Increased insulin signaling promotes faster proliferation and growth independent of changes to the distribution of cell cycle phases **A-B.** Protein synthesis visualized by OPP incorporation in control **(A)** and *InR-DA*-expressing **(B)** discs. Yellow dashed lines indicate the boundary between the wing pouch and hinge regions. The area above the line corresponds to the *rn-GAL4* expression domain, while the area below represents wild-type cells. The *InR-DA*-expressing wing pouch exhibits higher rates of protein synthesis, reflecting an increased biosynthetic capacity promoted by higher InR/PI3K/Akt activity. **C.** Quantification of mean OPP intensity in the pouch region of control and *InR-DA*-expressing discs. Mean and 95% CI is shown. Statistical significance was tested using two-tailed Unpaired t test, p value<0.0001. (control discs: n=5, *InR-DA-*expressing discs: n=6). **D-E**. EdU incorporation visualizes DNA replication in the pouch region of control **(D)** and *InR-DA*-expressing **(E)** discs. Dashed yellow squares highlight the magnified regions shown in **(D’,E’)**. *InR-DA*-expressing wing pouch shows higher levels of EdU intensities, reflecting accelerated speed of DNA replication. Rates of EdU incorporation in *InR-DA*-expressing discs were previously reported and quantified (Crucianelli et al., 2022). **F-G.** Cell cycle dynamics in control **(F)** and *InR-DA*-expressing **(G)** discs. FUCCI reporters *GFP-E2F1*^1-230^ (green) and *mRFP-NLS-CycB*^1-266^ (red) were used to visualize cell cycle phases. Note that expression of a constitutively active Insulin receptor does not alter the cell cycle phase profile of the wing pouch compared to control disc. Discs were stained with DAPI to visualize nuclei. Fluorescence intensities are reported as arbitrary units. Maximum projections of multiple confocal sections are shown in **F-G.** Sum projections of multiple confocal sections are shown in **A-B** and **D-E.** Scale bars: 100 μm in **A,B**, **D,E**, **F,G;** 50 μm in zoomed-in **D,E.**

**Figure S4.2.**
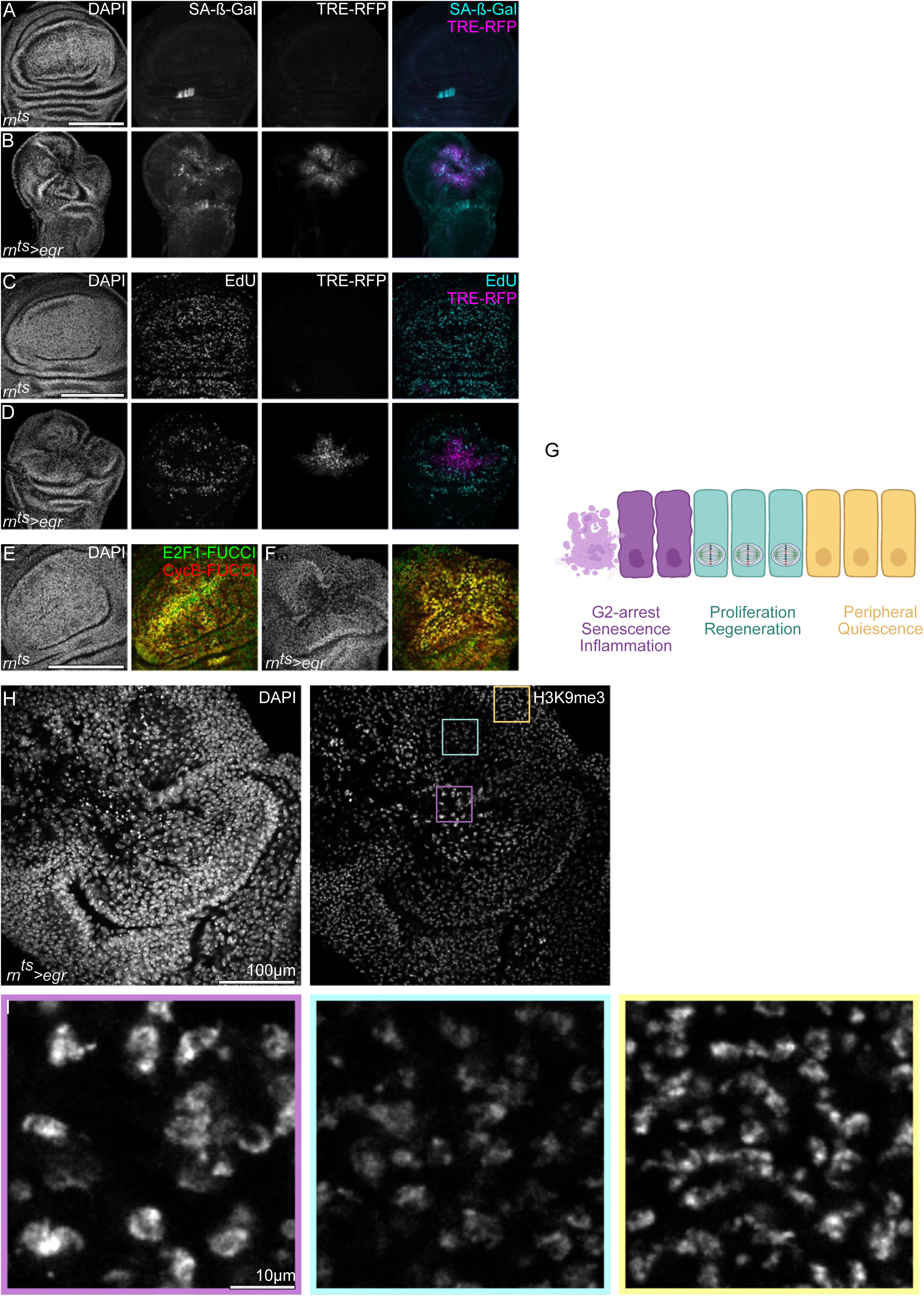
Persistence methyltransferase activity drives H3K9me3 accumulation through metabolic changes coupled to a senescent cell cycle arrest **A-B.** Senescence-associated β-galactosidase (SA-β-gal) activity (cyan or gray) in control **(A)** and *egr-* expressing **(B)** discs. TRE-RFP reporter (magenta) visualizes JNK-pathway activity. SA-β-gal activity in *egr-* expressing discs was previously reported and quantified (Jaiswal et al., 2023). **C-D. EdU incorporation (gray or cyan) visualizes DNA replication in control (B) and *egr-*expressing (D) discs. TRE-RFP reporter (magenta) visualizes JNK-pathway activity.** EdU incorporation was previously reported and quantified (Jaiswal et al., 2023, Floc’hlay et al., 2023). **E-F.** Cell cycle dynamics in control **(E)** and *egr*-expressing **(F)** discs. FUCCI reporters *GFP-E2F1*^1-230^ (green) and *mRFP-NLS-CycB*^1-266^ (red) were used to visualize cell cycle phases. **G.** A model illustrating inflammatory tissue damage in *Drosophila* wing imaginal discs. Eiger expression activates the JNK pathway, an early stress response, which induces a senescence-like cell-cycle arrest in the G2 phase at center of the tissue damage and triggers a senescence program (Cosolo et al., 2019). These arrested cells secrete Unpaired cytokines, which activate the JAK/STAT pathway in surrounding cells, promoting compensatory proliferation during tissue regeneration (Crucianelli et al., 2022; Floc’hlay et al., 2023; Herrera & Bach, 2019; Jaiswal et al., 2023; La Fortezza et al., 2016; Santabarbara-Ruiz et al., 2015; Worley et al., 2022). At the periphery of the damage site, cells remain in a quiescent state (Maya et al., 2024). **H-I.** Immunostaining for H3K9me3 in *egr*-expressing disc **(H).** The magenta frame highlights G2-arrested senescent cells, which show elevated H3K9me3 levels. The cyan frame marks cells undergoing compensatory proliferation, where H3K9me3 levels are reduced. The yellow frame indicates quiescent cells, which have higher H3K9me3 levels than proliferating cells **(I).** Discs were stained with DAPI to visualize nuclei. Maximum projections of multiple confocal sections are shown in **H.** Sum projections of multiple confocal sections are shown in **A-D** and **E-F.** Scale bars: 100 μm in **A-B**, **C-D**, **E-F** and **H**; 10 μm in **I**.

**Figure S5.1.**
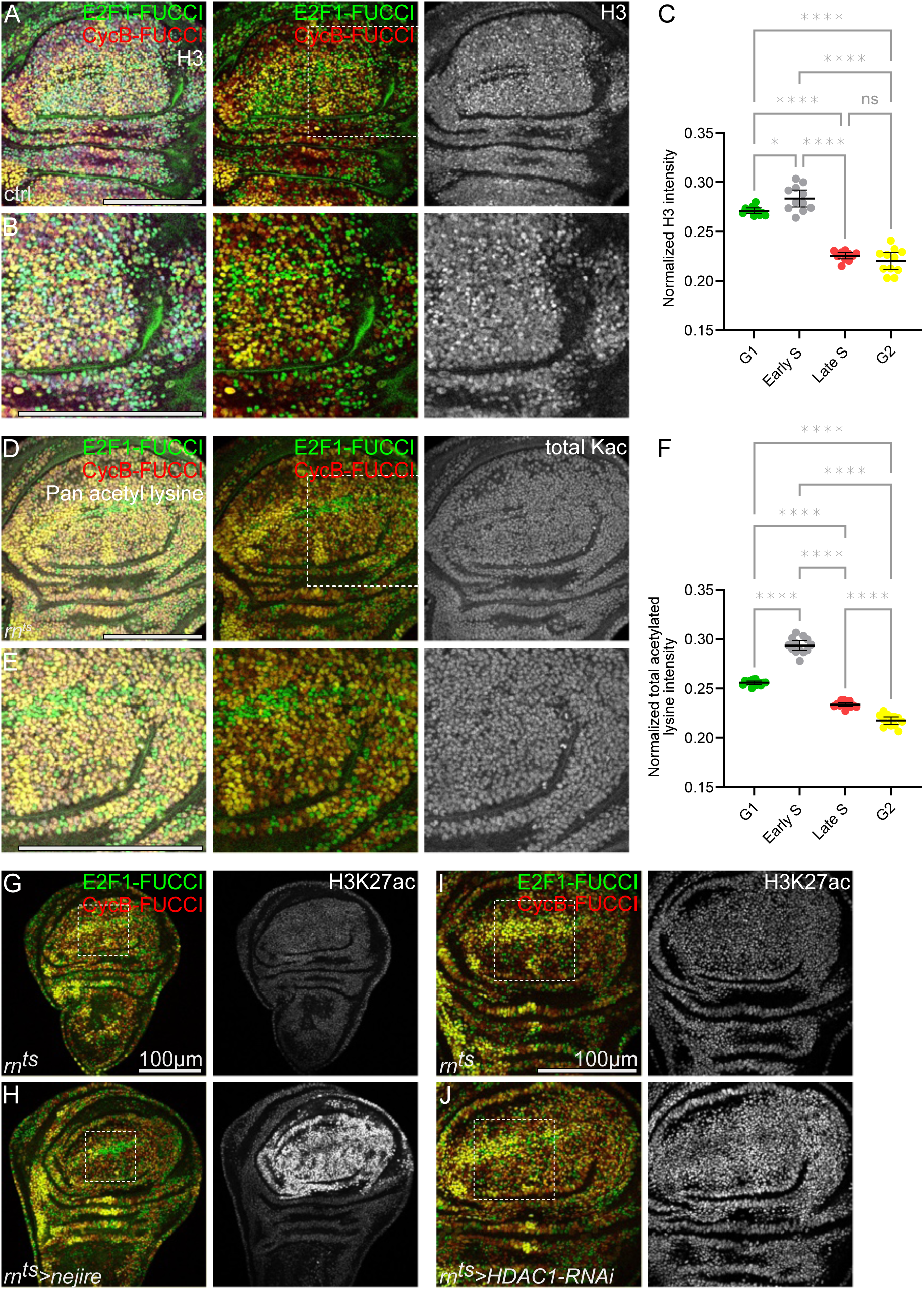
Total H3 and acetyl-lysine dynamics do not recapitulate the S phase-associated increase in H3K27ac **A-B.** Immunostaining for H3 in a developing control disc **(A**), with a magnified region (demarcated by a white dashed square) shown in panel B, also expressing the FUCCI reporter (*GFP-E2F1*^1-230^ (green) and *mRFP-NLS-CycB*^1-266^ (red)) to visualize cell cycle phases. **C.** Quantification of normalized H3 intensity level with respect to different cell cycle phases in the pouch region of control discs. Please note that normalization was performed against the average fluorescence intensity across all cell cycle phases. Mean and 95% CI is shown. Statistical significance was tested using Repeated Measures One-Way ANOVA followed by Tukey’s post-hoc test for multiple comparison (n=11 discs). Adjusted p= 0.0192 for G1 vs. Early S; p<0.0001 for G1 vs. Late S; p<0.0001 for G1 vs. G2; p<0.0001 for Early S vs. Late S; p<0.0001 for Early S vs G2 and p=0.4147 for Late S vs G2. **D-E.** Immunostaining for total acetylated lysine level in a developing control disc **(D),** with a magnified region (demarcated by a white dashed square) in panel E, also expressing the FUCCI reporters (*GFP-E2F1*^1-230^ (green) and *mRFP-NLS-CycB*^1-266^ (red)) to visualize cell cycle phases. **F.** Quantification of normalized total acetylated lysine level in different cell cycle phases in the pouch region of control discs. Please note that normalization was performed against the average fluorescence intensity across all cell cycle phases. Mean and 95% CI is shown. Statistical significance was tested using Repeated Measures One-Way ANOVA followed by Tukey’s post-hoc test for multiple comparison (n=13 discs). Adjusted p<0.0001 for G1 vs. Early S; p<0.0001 for G1 vs. Late S; p<0.0001 for G1 vs. G2; p<0.0001 for Early S vs. Late S; p<0.0001 for Early S vs G2 and p<0.0001 for Late S vs G2. **G-H.** Immunostaining for H3K27ac in control **(G)** and *nejire*-expressing **(H)** discs, also expressing the FUCCI reporters (*GFP-E2F1*^1-230^ (green) and *mRFP-NLS-CycB*^1-266^ (red)) to visualize cell cycle phases. The location of magnified regions shown in Main Figure 5F and G is indicated by white dashed squares. **I-J.** Immunostaining for H3K27ac in control **(I)** and *HDAC1-RNAi*-expressing **(J)** discs also expressing the FUCCI reporters (*GFP-E2F1*^1-230^ (green) and *mRFP-NLS-CycB*^1-266^ (red)) to visualize cell cycle phases. The location of magnified regions shown in Main Figure 5H and I is indicated by white dashed squares. Fluorescence intensities are reported as arbitrary units. Maximum projections of multiple confocal sections are shown in **D-E.** Scale bars: 100 μm.

**Figure S5.2.**
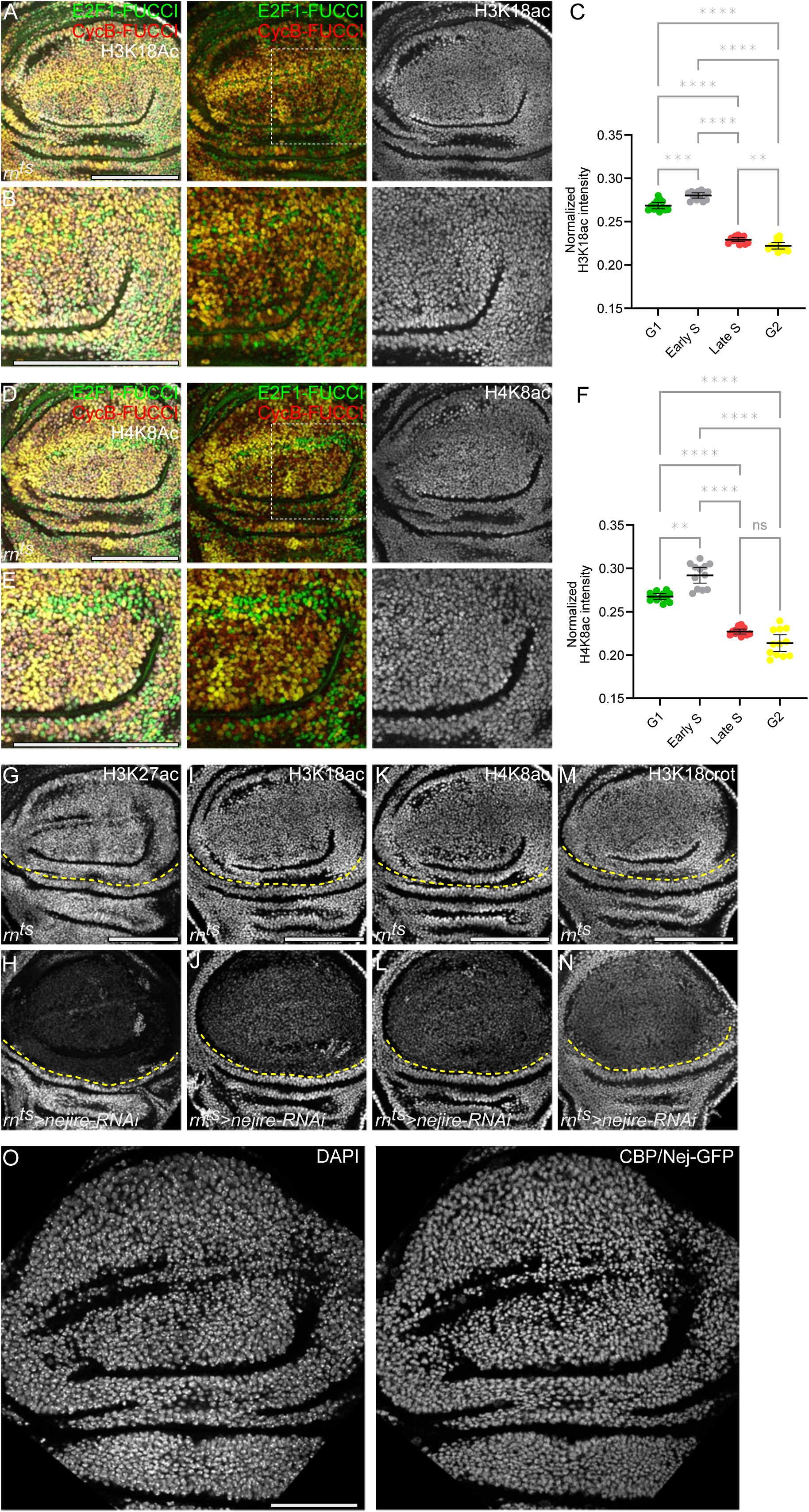
Total H3K18ac and H4K8ac dynamics only partially recapitulate the S phase-associated increase in H3K27ac **A-B.** Immunostaining for H3K18ac in a developing control disc **(A),** with a magnified region (demarcated by a white dashed square) in panel B, in the background of FUCCI reporters (*GFP-E2F1*^1-230^ (green) and *mRFP-NLS-CycB*^1-266^ (red)) to visualize cell cycle phases. **C.** Quantification of normalized H3K18ac intensity level in different cell cycle phases in the pouch region of control discs. Please note that normalization was performed against the average fluorescence intensity across all cell cycle phases. Mean and 95% CI is shown. Statistical significance was tested using Repeated Measures One-Way ANOVA followed by Tukey’s post-hoc test for multiple comparison (n=13 discs). Adjusted p=0.0003 for G1 vs. Early S; p<0.0001 for G1 vs. Late S; p<0.0001 for G1 vs. G2; p<0.0001 for Early S vs. Late S; p<0.0001 for Early S vs G2 and p=0.0042 for Late S vs G2. **D-E.** Immunostaining for H4K8ac in a normally developing control disc **(D),** with a magnified region (demarcated by a white dashed square) in panel E, also expressing the FUCCI reporters (*GFP-E2F1*^1-230^ (green) and *mRFP-NLS-CycB*^1-266^ (red)) to visualize cell cycle phases. **F.** Quantification of normalized H4K8ac intensity level in different cell cycle phases in the pouch region of control discs. Please note that normalization was performed against the average fluorescence intensity across all cell cycle phases. Mean and 95% CI is shown. Statistical significance was tested using Repeated Measures One-Way ANOVA followed by Tukey’s post-hoc test for multiple comparison (n=12 discs). Adjusted p=0.0018 for G1 vs. Early S; p<0.0001 for G1 vs. Late S; p<0.0001 for G1 vs. G2; p<0.0001 for Early S vs. Late S; p<0.0001 for Early S vs G2 and p=0.1076 for Late S vs G2. **G-H.** Immunostaining for H3K27ac in control **(G)** and *nejire-RNAi*-expressing **(H)** discs. Same disc is also shown in Figure 5D and E but repeated here to allow for direct comparison with **(I-N)**. **I-J.** Immunostaining for H3K18ac in control **(I)** and *nejire-RNAi*-expressing **(J)** discs. **K-L.** Immunostaining for H4K8ac in control **(K)** and *nejire-RNAi*-expressing **(L)** discs. **M-N.** Immunostaining for H3K18crot in control **(M)** and *nejire-RNAi*-expressing **(N)** discs. Yellow dashed lines indicate the boundary between the wing pouch and hinge regions. **O.** Expression of a Crisper-tagged CBP/nej-GFP (Marsh et al., 2025) in a developing disc. Discs were stained with DAPI to visualize nuclei. Fluorescence intensities are reported as arbitrary units. Maximum projections of multiple confocal sections are shown in **A-B** and **D-E.** Sum projections of multiple confocal sections are shown in **G-H, I-J, K-L** and **M-N.** Scale bars: 100 μm

**Figure S6.**
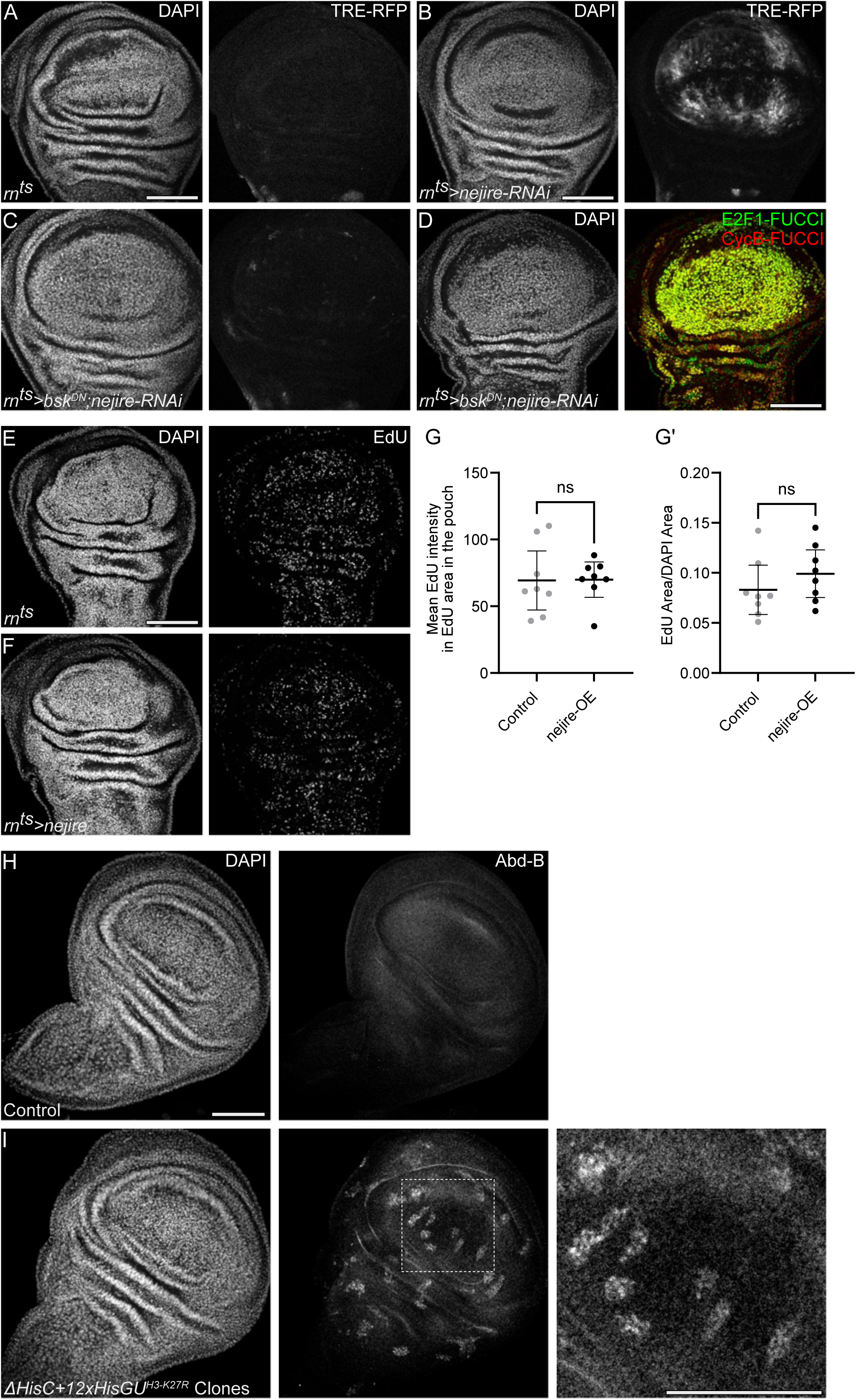
CBP/nej and H3K27ac are required to prevent replication stress **A-C.** Expression of the JNK pathway reporter TRE-RFP in control (A), *nejire-RNAi*-expressing (B) and *bsk^DN^, nejire-RNAi*-coexpressing (C) discs. **D.** Cell cycle dynamics in *bsk^DN^, nejire-RNAi*-coexpressing (D) disc. FUCCI reporters *GFP-E2F1*^1-230^ (green) and *mRFP-NLS-CycB*^1-266^ (red) were used to visualize cell cycle phases. Even when JNK pathway activity is suppressed, one can still observe a shift of cells to the G2 phase of the cell cycle *nejire-RNAi*-expressing discs. **E-F.** EdU incorporation to visualize DNA replication in the pouch region of control (E) and *nejire-* expressing (F) discs. **G.** Quantification of mean EdU intensity per EdU area in the pouch region of control and *nejire-* expressing discs, serving as a proxy for speed of DNA replication. Mean and 95% CI are shown. Statistical significance was tested using two-tailed Mann Whitney test, p value=0.5737 (control discs: n=8, *nejire-*expressing discs: n=8). **G’.** Quantification of EdU area per DAPI area in the pouch region of control and *nejire*-expressing discs, serving as a proxy number of cells undergoing DNA replication. Mean and 95% CI are shown. Statistical significance was tested using a two-tailed Mann Whitney test, p value=0.2345 (control discs: n=8, *nejire*-expressing discs: n=8). **H-I.** Immunostaining for Abd-B in control discs **(H)** and in discs expressing *12×His-GU^H3-K27R^* transgenes and containing clones homozygous mutant for a deletion of the histone gene cluster *ΔHisC* **(I)**. Discs were stained with DAPI to visualize nuclei. Fluorescence intensities are reported as arbitrary units. Sum projections of multiple confocal sections are shown in **A-C**, **E-F** and **H-I**. Scale bars: 100 μm.

---

**Table S1.** Experimental strains, reagents and software.

| Tool | Source | ID |
| --- | --- | --- |
| *w<sup>118</sup>* | David Bilder, University of California, Berkeley | FBal0018157 |
| *UAS-GFP<sup>S56T</sup>* | Bloomington *Drosophila* Stock Center | BDSC: 1521 |
| *rn<sup>GAL4-DeltaS</sup>, tubGAL80<sup>ts</sup>* | Bloomington *Drosophila* Stock Center | BDSC: 8142; 7018 recombinant |
| *rn<sup>GAL4-5</sup>, UAS-egr, tubP-GAL80<sup>ts</sup>* | Iswar Hariharan, University of California, Berkeley | BDSC: 8142; 7018 recombinant |
| '*E2F1-FUCCI, CycB-FUCCI*'; *Ubi-GFP.E2f1<sup>1-230</sup>, Ubi-mRFP1.NLS.CycB<sup>1-266</sup>* | Bloomington *Drosophila* Stock Center | BDSC: 55123 |
| *UAS-Cdk1 RNAi [TRiP.HMS01531]* | Bloomington *Drosophila* Stock Center | BDSC: 36117 |
| *lf/CyO-GFP; UAS-E2F, UAS-DP/TM6b* | Laura Buttitta, University of Michigan | PMID: 9657151 |
| *UAS-pnt.P1* | Bloomington *Drosophila* Stock Center | BDSC: 869 |
| *UAS-E(z)-RNAi (TRiP.HMS00066)* | Bloomington *Drosophila* Stock Center | BDSC: 33659 |
| *UAS-nejire-RNAi (TRiP.HMS01507)* | Bloomington *Drosophila* Stock Center | BDSC: 37489 |
| *UAS-InR-DA (UAS-InR.A1325D)* | Bloomington *Drosophila* Stock Center | BDSC: 8263 |
| '*TRE-RFP*': *TRE-DsRed.T4* | Dirk Bohmann, University of Rochester Medical Center | PMID: 22509270 |
| *UAS-nejire* | Bloomington *Drosophila* Stock Center | BDSC: 32573 |
| *UAS-HDAC1-RNAi (TRiP.HMS00607)* | Bloomington *Drosophila* Stock Center | BDSC: 33725 |
| *CBP-GFP* | Melissa Harrison, University of Wisconsin School of Medicine and Public Health | PMID: 40441155 |
| *w; Df(2L)HisC FRT40A/CyO, twi-GAL4, UAS-GFP; 6xHisGU<sup>H3K27R</sup>/6xHisGU<sup>H3K27R</sup>* | Jürg Müller, Max Planck Institute of Biochemistry | PMID: 23393264 |
| *yw hs-flp122; hs-nGFP FRT40A; 6xHisGU<sup>H3-K27R</sup>/SM5<sup>^</sup>TM6B* | Jürg Müller, Max Planck Institute of Biochemistry | PMID: 23393264 |
| *hs-flp122; FRT40A ubi-GFP/CyO (GFP); Dr/TM6c* |  |  |
| *UAS-bsk[DN]* | Bloomington *Drosophila* Stock Center | BDSC: 6409 |
| Chicken anti-GFP | Abcam | Cat. #: ab13970 |
| Rabbit monoclonal anti-GFP | Invitrogen | Cat. #: G10362 |
| Rat monoclonal anti-RFP | ChromoTek | Cat. #: 5F8 RRID: AB_2336064 |
| Rabbit polyclonal anti-H3K9ac | Active motif | Cat. #: 39017 RRID: AB_2616593 |
| Rabbit polyclonal anti-H4K8ac | Active motif | Cat. #: 61104 RRID: AB_2793506 |
| Rabbit polyclonal anti-acetyl Lysine | Abcam | Cat. #: ab80178 |
| Rabbit monoclonal anti-crotonyl Histone H3 | PTMab | Cat. #: PTM-517RM |
| Rabbit polyclonal anti-H3K9me1me2me3 | Active motif | Cat. #: 39242 RRID: AB_2793200 |
| Rabbit polyclonal anti-H3K18ac | Active motif | Cat. #: 39756 RRID: AB_2714186 |
| Mouse monoclonal anti-H3K27ac | Active motif | Cat. #: 39085 RRID: AB_2793305 |
| Rabbit polyclonal anti-H3K4me3 | Abcam | Cat. #: ab8580 |
| Mouse monoclonal anti-H3K27me3 | Abcam | Cat. #: ab6002 |
| Rabbit polyclonal anti-H3K9me3 | Abcam | Cat. #: ab8898 |
| Rabbit polyclonal anti-H3 | Abcam | Cat. #: ab1791 |
| Rabbit polyclonal anti-Histone H2AvD phosphoS137 | Rockland | Cat. #: 600-401-914 |
| Mouse monoclonal anti-Cut | DSHB | Cat. #: 2B10 RRID: AB_528186 |
| Mouse monoclonal anti-Engrailed/Invected | DSHB | Cat. #: 4D9 RRID: AB_528224 |
| Mouse monoclonal anti-Nubbin | DSHB | Cat. #: 2D4<br>RRID: AB_2722119 |
| Mouse monoclonal anti-Antp | DSHB | Cat. #: 8C11<br>RRID: AB_528083 |
| Mouse monoclonal anti-Ptc | DSHB | Cat. #: Apa1<br>RRID: AB_528441 |
| Mouse monoclonal anti Abd-B | DSHB | Cat. #: 1A2E9<br>RRID: AB_528061 |
| Mouse monoclonal anti-Wg | DSHB | Cat. #: 4D4<br>RRID: AB_528512 |
| Mouse monoclonal anti-Histone H3<br>(for CUT&Tag) | Active motif | Cat. #: 39763<br>RRID: AB_2650522 |
| Rabbit polyclonal anti-H3K27ac (for<br>CUT&Tag) | Diagenode | Cat. #: C15410196 |
| Rabbit polyclonal anti-H3K27me3 (for<br>CUT&Tag) | Diagendoe | Cat. #: C15410195 |
| Rabbit monoclonal anti-H3K9me3 (for<br>CUT&Tag) | Abcam | Cat. #: ab176916 |
| DAPI | Sigma Aldrich | Cat #: D9564 |
| Goat anti-mouse IgG Alexa Flour 488 | Invitrogen | Cat. #: A-11001 |
| Goat anti-chicken IgY Alexa Flour 488 | Invitrogen | Cat. #: A-11039 |
| Goat anti-rabbit IgG Alexa Flour 488 | Invitrogen | Cat. #: A-11008 |
| Goat anti-mouse IgG Alexa Flour 555 | Invitrogen | Cat. #: A-21422 |
| Goat anti-rat IgG Alexa Flour 555 | Invitrogen | Cat. #: A-21434 |
| Goat anti-mouse Alexa Flour 647 | Invitrogen | Cat. #: A-21235 |
| Donkey anti-mouse Alexa Flour 647<br>preabsorbed | Abcam | Cat. #: ab150111 |
| Goat anti-rat Alexa Flour 647 | Invitrogen | Cat. #: A-21247 |
| Goat anti-rabbit Alexa Flour 647 | Invitrogen | Cat. #: A-21244 |
| Click-iT Plus EdU Alexa Fluor 647<br>Imaging Kit | Invitrogen | Cat. #: C10640 |
| Click-iT Plus OPP Alexa Fluor 647<br>Imaging Kit | Invitrogen | Cat. #: C10458 |
| CellEvent Senescence Green<br>Detection Kit | Invitrogen | Cat. #: C10850 |
| FIJI (ImageJ 1.54f) | (Schindelin et al., 2012) | https://fiji.sc/ |
| GraphPad Prism | Graph Pad | version 9.5.0 |
| Affinity Designer | Affinity | Up to version 2.6.3 |
| Biorender | Science Suite Inc. DBA<br>BioRender | Biorender |

